# Semantic reasoning takes place largely outside the language network

**DOI:** 10.64898/2025.12.07.692873

**Authors:** Anna A. Ivanova, Carina Kauf, Ruimin Gao, Jingyuan Selena She, Hope H. Kean, Tanya Goldhaber, Alfonso Nieto-Castañón, Rosemary Varley, Nancy Kanwisher, Evelina Fedorenko

**Author notes:** Co-senior authors (Nancy Kanwisher, Evelina Fedorenko). **Author Contributions**: https://docs.google.com/spreadsheets/d/16NyApGgAl8B7e664p4Ix1BHNEWKU3uPGtnjCTUB5BvE/edit?usp=sharing.

## Abstract

The brain’s language network is often implicated in the representation and manipulation of abstract semantic knowledge. However, this view is inconsistent with a large body of evidence suggesting that language processing is neurally distinct from the rest of cognition. Here, we use precision brain imaging to uncover a set of brain regions, separate from the language network, that are engaged in semantic processing of both linguistic and pictorial stimuli. In three fMRI experiments, participants (total n=41 tested across 49 sessions) viewed sentences and pictures depicting simple events. In separate blocks, they performed either a semantic task or a difficulty-matched perceptual task. Across all three experiments, several areas in left lateral prefrontal cortex, left temporo-parietal cortex, and right cerebellum responded to semantic tasks for both sentences and pictures. These semantic processing areas are spatially and functionally distinct from the nearby language-selective areas, as well as from the multiple demand and default mode networks, exhibiting a unique response profile. Our results provide evidence for a new kind of selectivity in the human brain and pave the way for future explorations of the neural mechanisms that underlie semantic reasoning.

## Introduction

The capacity for abstract thought is a hallmark of intelligence. We humans constantly generalize across scenes, sounds, and sensations to extract common conceptual representations that can be widely used in a range of future situations. For instance, suppose you have a cat called Tibbles. Without a concept for Tibbles, you would need separate links among (i) the image of Tibbles, (ii) the softness of his fur, (iii) the sound of his meows, and (iv) his name. But with the abstract concept of Tibbles that spans these perceptual features, we need to only link each feature to that concept. Moreover, the concept of “Tibbles” is then connected to other concepts, like “cat”, “animal”, “likes milk”, and “purrs when patted”, giving us a powerful engine for inferring what Tibbles might be like and how to best care for him. What are the neurocognitive systems that support the human ability to represent and manipulate abstract information that generalizes across the surface form of the input?

One classic idea is that language serves as a vehicle for conceptual thought (Carruthers, 2002; Chomsky, 2007; Hinzen, 2013; Vygotsky, 1934). According to this view, linguistic representations are essential for organizing and reasoning over conceptual knowledge. In apparent accordance with this theory, a number of brain imaging studies have reported overlapping activations for words/sentences and pictures in left-lateralized frontal and/or temporal/temporo-parietal brain regions (e.g., Devereux et al., 2013; Fairhall & Caramazza, 2013; Handjaras et al., 2017; Krieger-Redwood et al., 2015; Shinkareva et al., 2011; Vandenberghe et al., 1996; Visser et al., 2012; Wurm & Caramazza, 2019). The anatomical location of these areas broadly resembles the human language network, a set of regions that respond consistently and selectively to language (Fedorenko et al., 2024). A parallel line of research has argued that a set of frontal and temporal regions —again, resembling the language network—carry out so-called “semantic control”, i.e., the selection of relevant semantic information in response to contextual/task demands (e.g., Davey et al., 2016; Jackson, 2021; Jackson et al., 2021; Jefferies, 2013; Lambon Ralph et al., 2017; Martin & Chao, 2001; Whitney et al., 2011, 2012; M. Zhang et al., 2022).

A straightforward interpretation of the neuroscience evidence to date is that language and semantic reasoning share the same neural mechanisms. Such a claim would rely on reverse inference from anatomy (Poldrack, 2006, 2011). But frontal and temporal cortices—where semantic activation is commonly observed—are highly functionally heterogeneous, with nearby regions showing distinct profiles (Braga et al., 2019; Deen et al., 2015; Du et al., 2024; Fedorenko & Blank, 2020; for a review, see Gratton & Braga, 2025). Thus, the fact that a semantic task engages a left frontal area or a left temporal area does not automatically mean that these areas are the same as the areas that support language processing. Furthermore, many studies of semantic processing rely solely on linguistic stimuli, making it impossible to disentangle language comprehension and semantic reasoning. To test the relationship between the brain areas underlying these two processes, we need (i) a precision brain imaging approach, which localizes functional brain areas in individual participants, and (ii) an experimental condition where participants perform semantic reasoning over both linguistic and non-linguistic inputs, such as pictures.

Evidence that the language system may *not* be the primary locus of abstract semantic processing comes from both behavioral and neural investigations. Potter and Faulconer (1975) found that semantic information could be extracted from pictures as fast or faster than the verbal labels of the pictured objects, suggesting that conceptual information need not be routed through language. More recently, we have shown that the language system responds strongly to sentences and much less strongly to pictures and movies (Benn, Ivanova et al, 2023; Ivanova et al., 2021; Shain, Paunov, Chen et al., 2023; Sueoka et al., 2024); and some patients with severe aphasia can perform semantic tasks on visual inputs (e.g., Ivanova et al., 2021; Varley et al., 2001). Moreover, pre-verbal infants (Hirsh-Pasek & Golinkoff, 2010; Spelke, 2022) and many non-human animals (Beran et al., 2014; Bräuer et al., 2020; Smith et al., 2016) show a remarkable capacity for conceptual abstraction across physical and social domains, suggesting that semantic processing, too, may take place in the absence of language.

These findings raise the possibility that semantic reasoning need not rely on linguistic representations. On this view, the language system extracts relatively shallow semantic information from words and sentences, but considerable further elaboration outside the language system is required to build the kind of semantic representations that form the stuff of thought.

The goal of this study is to explore the neural basis of semantic reasoning. We define semantic reasoning as the process of accessing and operating over semantic knowledge to achieve a specific goal (e.g., answer a question). Two key factors allow us to systematically compare semantic reasoning and linguistic processing. First, we examine semantic reasoning over both linguistic stimuli (sentences) and non-linguistic stimuli (pictures). A putative semantic reasoning brain region/network should respond strongly during semantic processing regardless of input format, showing no preference for sentences over pictures. Second, we functionally localize language and semantic regions in individual participants (Fedorenko et al., 2010), an approach that overcomes the limitations of reverse inference and enables us to determine the true extent of the language network’s involvement in semantic reasoning.

We consider four nonexclusive possibilities for the loci for semantic reasoning: (1) a specialized set of brain regions that responds to inputs across modalities and is functionally selective for semantic reasoning; (2) the language network, commonly associated with semantic cognition (Binder et al., 2009); (3) the multiple demand network (Duncan, 2010; Duncan & Owen, 2000), also known as the fronto-parietal network, which is active during many different cognitively challenging tasks (Assem et al., 2020; Fedorenko et al., 2013; Hugdahl et al., 2015; Shashidhara, Mitchell, et al., 2019); and (4) the default mode network (DMN; Baldassano et al., 2018; Z. Hu et al., 2019; Jouen et al., 2015; cf. Thierry & Price, 2006), which has been hypothesized to support construction of structured mental situation models (Baldassano et al., 2017; Hassabis & Maguire, 2009) and has been implicated specifically in semantic reasoning (Fernandino & Binder, 2024; Jefferies & Smallwood, 2025). We also consider a possibility that semantic cognition is embodied (Barsalou, 2008; Pulvermüller & Fadiga, 2010), in which case we would either observe strong responses to both linguistic and non-linguistic in sensorimotor brain regions or fail to find crossmodal semantic regions at all (which would mean that all brain areas show a modality preference).

To foreshadow our findings, we identified a set of left-lateralized cortical and right-lateralized cerebellar brain areas that respond to both sentence and picture meanings, and are distinct from the language network, as well as from the multiple demand and default mode networks. Thus, we conclude that semantic reasoning recruits its own neural machinery.

## Results

Across three experiments, we probe neural responses to sentences and pictures to uncover brain areas that support amodal semantic processing. All three experiments followed the same design (**Figure 1A**)—crossing stimulus type (sentences, pictures) and task (semantic, perceptual), approximately matched for difficulty (see **SI-1** for behavioral performance metrics). The design variations across the three experiments enable us to ensure the generalizability of our results across task (plausibility vs. action reversibility judgments), event type (animate-inanimate or animate-animate interactions), and picture type (real photos vs. line drawings). Data from a total of 41 unique individuals are included in the analyses (21 women, mean age = 29, SD = 9.1; n=12 in Experiment 1, n=15 in Experiment 2, and n=22 in Experiment 3; 7 participants overlapped between Experiments 2 and 3; see <u>Methods</u> for further details on the experimental design).

**Figure 1.**
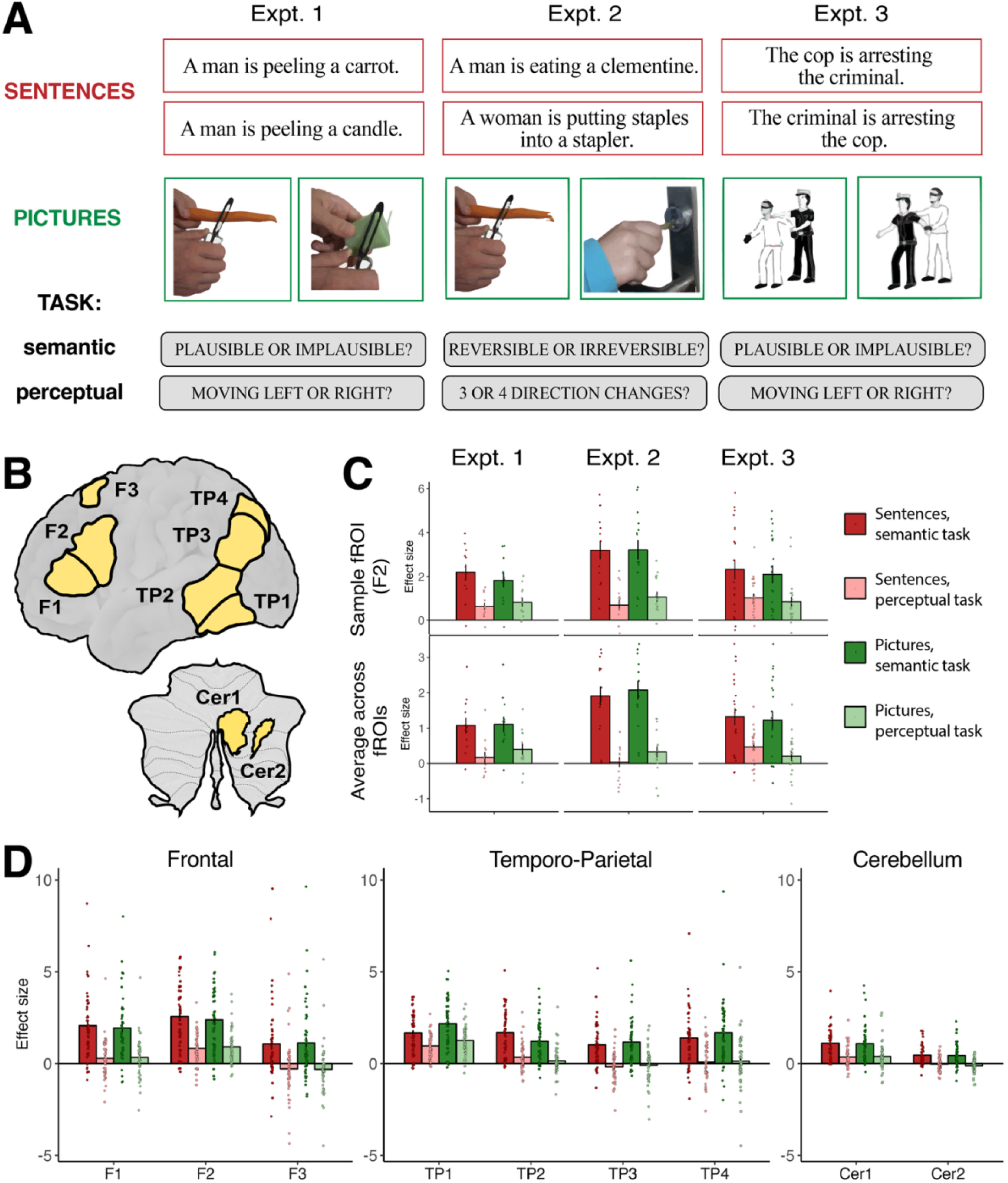
A set of brain regions responds during semantic reasoning over pictures and sentences. **(A)** Experimental design. The three experiments all include a semantic and a perceptual task over sentences and pictures, but the exact stimuli and tasks vary, enabling tests of generalizability across experimental design variations. **(B)** Parcels marking the approximate anatomical locations of semantic brain regions. Individual-level semantic fROIs constitute 10% of the parcel volume; see Figures 2B and **3B** for sample individual-specific fROIs. **(C)** Responses of the semantic fROIs are consistent across the three experiments and are similar between sentences and pictures. **(D**) Response profiles are similar across fROIs: all show selectivity for semantic vs. perceptual tasks across stimulus types. For experiment-wise fROI responses, see **Figure S2**. Here and elsewhere, each dot is a participant, and errorbars denote the standard error of the mean.

A key distinguishing feature of our work compared to previous investigations of semantic processing of linguistic vs. visual is the use of precision fMRI, where all analyses are performed within individuals, instead of averaging neural responses in each voxel across participants stimuli (cf. Popham et al., 2021). Because the association cortex is highly heterogeneous and the precise locations of functional areas vary across individuals (Blank et al., 2017; Fedorenko & Kanwisher, 2009; Frost & Goebel, 2012; Shashidhara, Spronkers, et al., 2019; Tahmasebi et al., 2012; Vázquez-Rodríguez et al., 2019), the group-averaging analyses are bound to overestimate overlap between nearby distinct areas (Nieto-Castañón & Fedorenko, 2012). This blurring of activations can lead to a false appearance that a single region contributes to two distinct functions (the alternative is that these functions are carried out by two distinct but adjacent areas). It can also conflate putative semantic reasoning regions with nearby cortical areas, including, critically, the language network (Fedorenko et al., 2024), but also the multiple demand network (Duncan et al., 2022) and the default mode network (Buckner & DiNicola, 2019), all of which have been implicated in aspects of semantic reasoning.

### Brain regions in the left frontal cortex, left temporo-parietal cortex, and right cerebellum respond during semantic reasoning over both sentences and pictures

We leveraged the similarity of the design across the three experiments and used the combined dataset to search across the brain for regions (“parcels”) that show an amodal semantic response signature—a stronger response to the semantic than the perceptual task for both sentences and pictures (see <u>Methods</u>). We identified nine such parcels (**Figure 1B**): three in left frontal cortex (F1-F3), four in left posterior temporal and inferior parietal cortex (TP1-TP4), and two in the right cerebellum (Cer1 and Cer2). Two additional regions were identified in the bilateral primary visual cortex, plausibly because of the higher visual processing demands in the semantic task; they are not included in the main analyses but are described in **SI (Figure S3)**. For subsequent analyses, we define participant-specific semantic functional regions of interest (*semantic fROIs*) within these parcels and examine their response profiles using independent data.

Statistical analyses of response magnitudes in the individually defined semantic fROIs, estimated in independent data (<u>Methods</u>), showed a significantly higher response to the semantic than perceptual tasks (network-level effect: β=1.19, p<0.001), with no main effect of stimulus type (sentences vs. pictures) and no interaction between task and stimulus type. Further, the response to the semantic conditions was above baseline (β=1.48, p<0.001). This amodal semantic response was remarkably consistent across the three experiments, which further establishes generalizability across stimuli, event types, and semantic tasks (**Figure 1C**). Specifically, follow-up analyses confirmed the main effect of semantic>perceptual task, no effect of stimulus type, and no interaction in each of the three experiments individually. This response profile was also consistent in individual fROIs (**Figure 1D**): all showed stronger responses to semantic than perceptual tasks (ps<0.001), above-baseline semantic task responses (ps<0.001), and no interaction between task and stimulus type. Most fROIs responded similarly strongly to sentences and pictures, except for TP1, which showed a preference for pictures (β=-0.81, p<0.001), and TP2, which showed a preference for sentences (β=0.57, p=0.03). The semantic regions’ response profile was robust, generalizing to a group of participants not used for parcel definition (**Figure S4**). Overall, we find several brain regions that exhibit strong selectivity for semantic over perceptual tasks across stimulus type.

### The semantic regions are largely distinct from the language network

The general topography of the amodal semantic regions bears some similarity to the canonical language network, which is also left-lateralized and has key components in the inferior frontal and posterior temporal / temporo-parietal cortex (Fedorenko et al, 2024), with additional components in the right cerebellum (Casto, Small, et al., 2025; LeBel & D’Mello, 2023; Wolna et al., 2025). So, next, we systematically examined the relationship between the amodal semantic regions and the language areas, identified with a standard localizer task (Fedorenko et al., 2010).

First, we tested whether the language network is the site of amodal semantic processing by examining its responses to the conditions of the critical experiments. We find that the language areas show a semantic>perceptual effect (β=0.76, p<.001), but in contrast to the amodal semantic areas, they also show a strong effect of stimulus type, with a preference for sentences over pictures (β=0.86, p<.001), as well as an interaction between task and stimulus type (β=0.52, p<.001). This result was previously reported for Experiment 3 in Ivanova et al. (2021; see Sueoka et al., 2024 for concordant findings using naturalistic stimuli), and replicates here across the full set of data from the three experiments (**Figure 2A**). Further, these areas show a robust preference for sentences over nonwords in the language localizer experiment, as estimated in independent data (β=1.35, p<.001) in which participants perform no task over the sentences they view. Thus, the core language regions show a strong preference for sentences.

**Figure 2.**
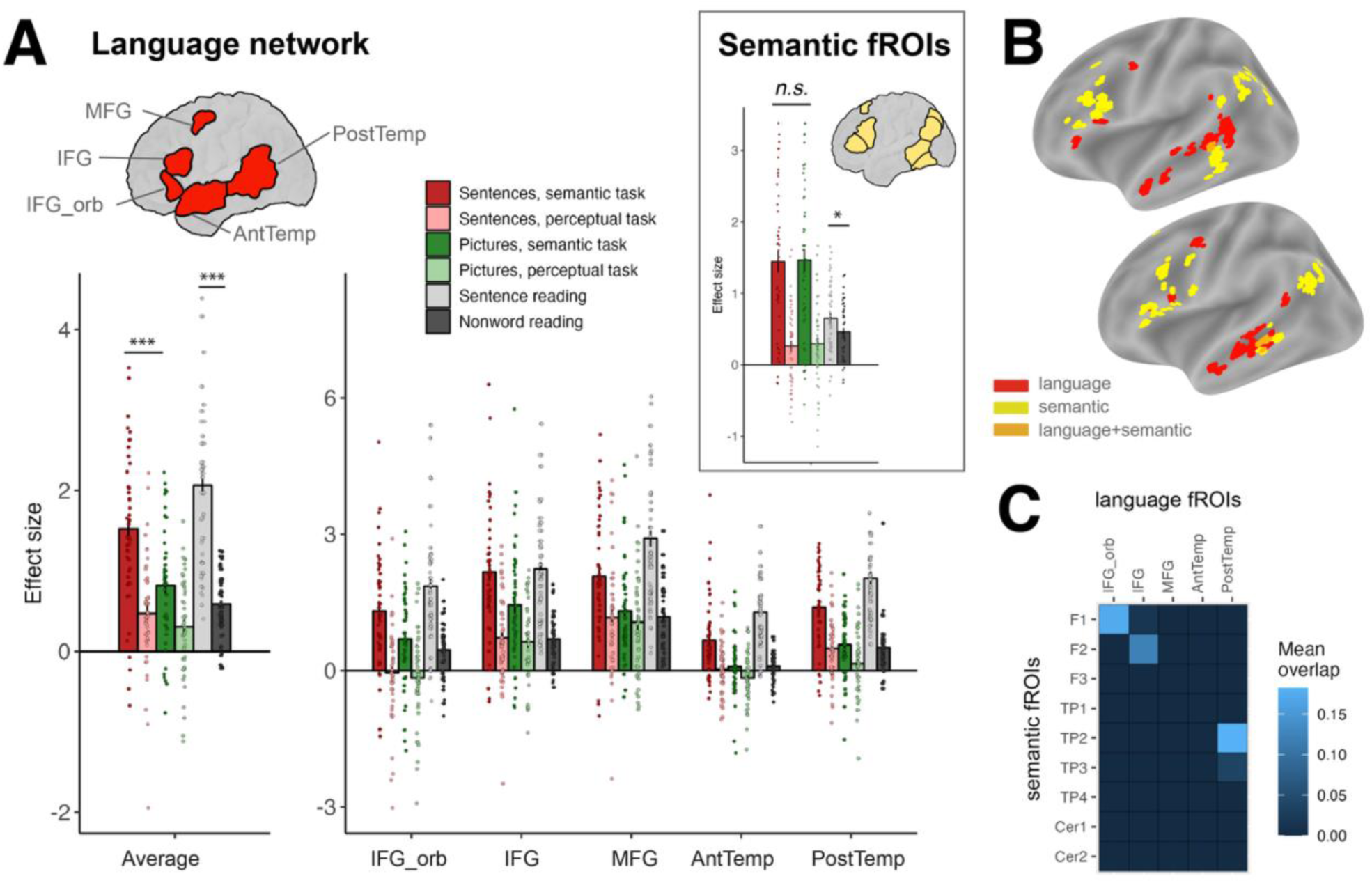
Semantic regions are distinct from the language network. **(A)** The response profile of the language network (left: average, right: individual fROIs) to critical task conditions (semantic and perceptual tasks over sentences and pictures) and the language localizer conditions (sentence reading and nonword reading), estimated in independent data. **Inset**: average response of the semantic fROIs to the same conditions. **(B)** Language and semantic fROIs in two individual participants. **(C)** Overlap coefficients between semantic and language fROIs, averaged across participants.

Second, we examined the responses in the amodal semantic regions to the language localizer and found that only two fROIs—both in the temporal lobe—exhibited a significant sentences>nonwords effect: TP2 (β=0.72, p<.001) and TP3 (β=0.52, p=.009), with others exhibiting equally low responses to sentence and nonword reading (with no task). All semantic fROIs—including TP2 and TP3—showed a significantly stronger response to the semantic task on sentences vs. passive sentence reading (ps<0.001), indicating that they are primarily driven by the semantic task, not simply reading and understanding sentences.

Finally, we directly examined the spatial overlap between language and semantic fROIs in individual participants’ brains (**Figure 2B, 2C**). We find modest overlap between language and semantic fROIs in posterior temporal lobe (TP2 vs. the PostTemp language fROI) and in the inferior frontal gyrus (F1 vs. the IFG_orb fROI; and F2 vs. the IFG fROI), but overall, the two systems are spatially largely separate, with nearby but non-overlapping spatial locations.

Together, our comparisons of language and semantic areas indicate that these regions are adjacent but largely distinct, with different functional response profiles and spatial locations. The key distinction is that the language areas prefer sentences to pictures, whereas the semantic areas engage in semantic task processing regardless of stimulus modality.

### The semantic regions are largely distinct from the multiple demand and default mode networks

We also compared the response profile and spatial locations of semantic fROIs with two other known networks: multiple demand, implicated in domain-general task-driven processing (Duncan, 2010; Fedorenko et al., 2013), and default mode network, sometimes implicated in semantic processing specifically (Fernandino & Binder, 2024; Jefferies & Smallwood, 2025). These two networks can be identified functionally with a demanding executive function task (e.g., a working memory task), using a hard>easy contrast for the multiple demand network (Fedorenko et al., 2013; Shashidhara et al., 2019), and the reverse contrast for the default mode network (e.g., Mineroff, Blank et al., 2018).

We found that both networks exhibit a distinct response profile from the semantic fROIs (**Figure 3A**). The multiple demand network exhibits a classic difficulty-driven response, with a hard spatial working memory condition evoking more activation than an easier spatial working memory condition, as estimated in independent data (LH: β=1.19, p<.001, RH: β=1.46, p<.001). The default mode network shows the opposite pattern, with lower responses to the harder spatial working memory condition relative to easy spatial working memory (LH: β=-0.76, p<.001, RH: β=-0.66, p<.001). The semantic areas, however, showed no difference between the hard and the easy spatial working memory conditions (β=0.06, *n.s.*) and an overall low response, meaning that these areas are neither activated nor deactivated by the difficulty of a non-semantic cognitive task.

**Figure 3.**
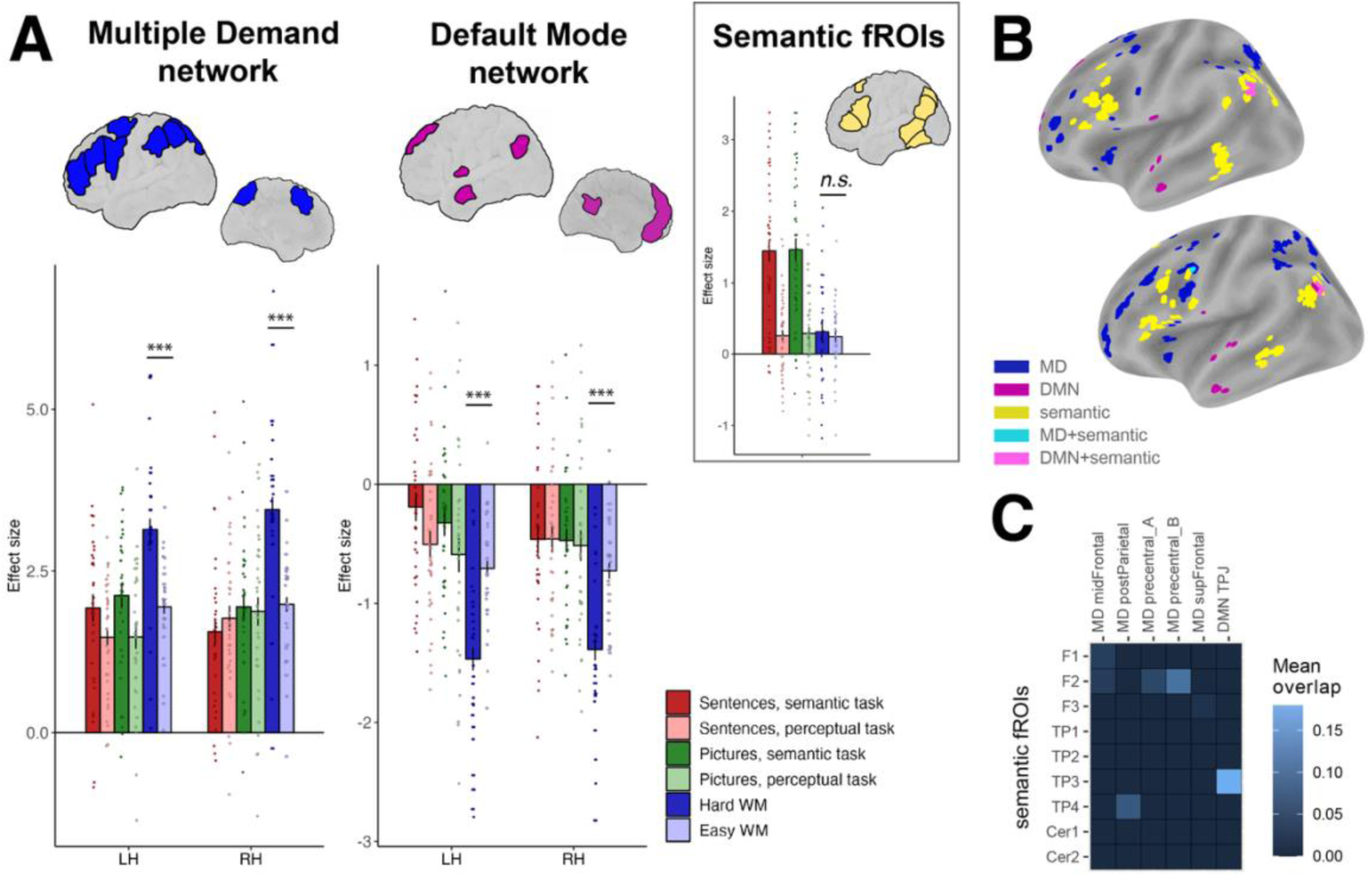
Semantic regions are distinct from the multiple demand and default mode networks. **(A)** The response profile of the multiple demand network (left) and the default mode network (right) to critical task conditions (semantic and perceptual tasks over sentences and pictures) and a spatial working memory (WM) task conditions (that serve to localize the multiple demand and default mode networks). **Inset**: average response of the semantic fROIs to the same conditions. **(B)** Multiple demand, default mode network, and semantic fROIs in two individual participants. MD=multiple demand, DMN=default mode network. **(C)** Overlap coefficients between semantic and multiple demand/default mode network fROIs, averaged across participants. Only multiple demand and default mode network fROIs with overlap above 0.01 are shown.

We also investigated the extent to which multiple demand and default mode networks contribute to semantic processing. In the multiple demand network, the response to the semantic task conditions was above baseline (LH: β=2.16, p<.001, RH: β=1.78, p<.001). Further, the left hemisphere multiple demand fROIs showed a semantic>perceptual effect, but not the right hemisphere (LH: β=0.58, p<.001, RH: β=-0.09, *n.s.*). Therefore, the multiple demand fROIs are driven by both semantic and perceptual tasks, with left hemisphere fROIs showing a slight preference for the semantic task. The default mode network showed below-baseline responses to the semantic task (LH: β=-0.27, p=.02, RH: β=-0.49, p<.001). The left but not the right hemisphere default mode network fROIs showed a small semantic>perceptual effect (LH: β=0.28, p=.02, RH: β=0.0, *n.s.*). Therefore, the left-hemisphere default mode network areas do show a slight preference for the semantic task, but responses fall below baseline, in contrast to the robust above-baseline response in the semantic areas.

Finally, an overlap analysis of semantic fROIs vs. multiple demand and default mode network fROIs (**Figure 3B, 3C**) confirmed that these systems are largely non-overlapping. The strongest point of overlap was between TP3 and default mode network left TPJ/angular gyrus. We hypothesize that this area may serve as a linking point between semantic and default mode network areas (Xu et al., 2016) but leave further evaluation of this hypothesis to future work.

### Contralateral homotopes of the semantic regions show a preference for pictures over sentences

The semantic regions we found are markedly lateralized, with all parcels falling either in the left cerebral hemisphere or in the right cerebellum (right cerebellar regions connect to the left neocortical regions; Buckner et al., 2011a). Does it mean that the contralateral regions are not engaged in semantic processing, or perhaps those regions were just not robust enough to emerge in a whole-brain search (similar to the vATL regions)? To find out, we mirror-projected the semantic parcels onto the opposite hemisphere and repeated our fROI analysis (**Figure S5**).

All contralateral fROIs show above-baseline responses to the semantic task (most ps<.001, Cer2: p=.002, F3: p=.033) and stronger responses to the semantic task relative to the perceptual task (max p=.008). However, unlike the original set of LH cortical and RH cerebellar semantic regions, most contralateral fROIs also show a significant stimulus type effect, with stronger responses to pictures than sentences (except F3 and Cer2, which had low responses overall). No regions showed an interaction between task and stimulus type.

Multiple contralateral fROIs showed a numerical hard>easy effect in the spatial working memory task, although it only reached significance in F2 (β=.57, p=.036). No fROI showed a difference in response to sentence reading vs. nonword reading, and all except three (TP1, TP3, and Cer2) showed stronger responses to sentences during a semantic task than during passive reading.

Overall, the fROIs defined within the homotopic counterparts of semantic parcels show sensitivity to semantic processing, but they show a preference toward pictorial over sentence stimuli.

### Bilateral regions in ventral anterior temporal lobe respond during both semantic reasoning and passive sentence comprehension

The ventral anterior temporal lobe (vATL) figures prominently in theories of semantic cognition (Patterson et al, 2007; Lambon Ralph et al, 2017), yet our semantic parcels did not include any anterior temporal regions. To determine whether this absence was caused by signal dropout, an issue that commonly affects this region due to its proximity to the ear cavity (Halai et al., 2014), we computed tSNR—a measure of signal quality (Welvaert & Rosseel, 2013)—in the semantic parcels and ATL parcels defined using the Desikan (2006) anatomical atlas (see **SI** text and **Figure S6** for details). A two-sample t-test revealed significantly higher tSNR values within the semantic parcels compared to the anatomical ATL parcels (t = 57.6, p < 0.001). However, the average tSNR in the ATL parcels was 55, substantially higher than the threshold of 20, which has been previously suggested as a minimum criterion for examining ATL responses (Binder et al., 2011). Thus, we conducted a follow-up analysis to specifically examine responses in semantic fROIs defined within the ATL parcels.

We found that, for our main task, the vATL fROIs showed a similar response profile to the frontal, temporo-parietal, and cerebellar semantic regions discovered in our whole-brain search (**Figure 4A**). Specifically, they showed a main effect of task (Semantic>Perceptual, left vATL: β=0.78, p<.001, right vATL: β=0.53, p<.001), no main effect of stimulus type, and no interaction between task and stimulus type. However, unlike most semantic fROIs in the main set, vATL fROIs also showed a significant effect for sentence reading > nonword reading (left vATL: β=0.34, p<.001, right vATL: β=0.18, p=.011), indicating that they respond to semantically meaningful stimuli even in the absence of a semantic task (see Wolna et al., 2025, for concordant evidence from a large-scale dataset). In fact, there was no difference between the sentence semantic task and the passive sentence reading conditions (left vATL: β=-0.06, *n.s*., right vATL: β=-0.07, *n.s.*).

**Figure 4.**
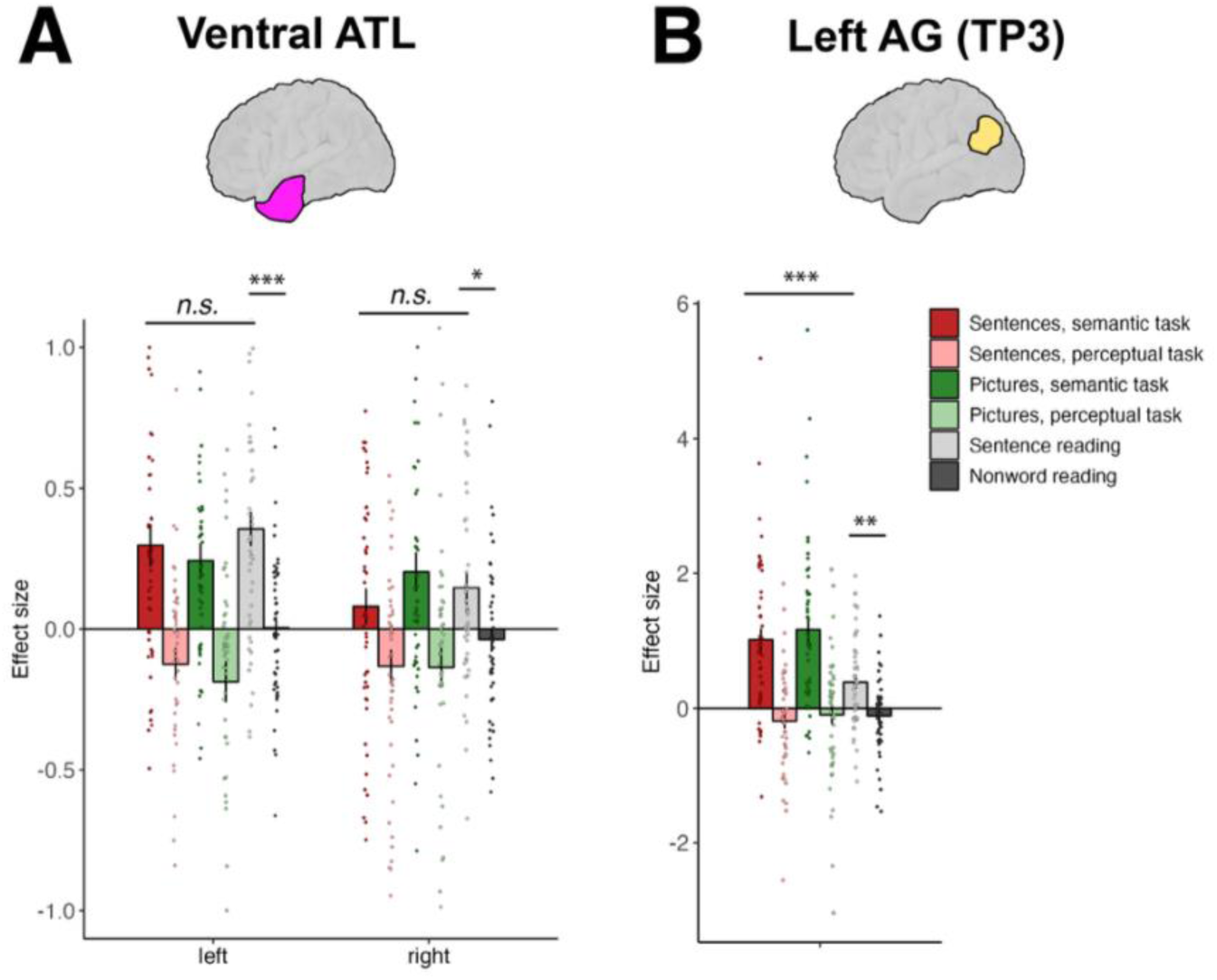
The response profiles of semantic fROIs in brain regions commonly implicated in semantic processing: **(A)** left and right ventral anterior temporal lobes (vATL), which did not emerge in our whole-brain analysis (potentially due to lower tSNR in these areas), and **(B)** left angular gyrus (AG), identified in our whole-brain analysis as parcel TP3. The plots show responses to critical task conditions (semantic and perceptual tasks over sentences and pictures) and to the language localizer conditions (sentence reading and nonword reading).

It is informative to directly compare the response profile of vATL with the response profile of our semantic region TP3, which falls mainly within the left angular gyrus (AG)—another anatomical area commonly implicated in semantic processing (Graves et al., 2023; Kuhnke, Chapman, et al., 2023; Seghier, 2013) (**Figure 4B**). As mentioned previously, this area shows a main effect of task (Semantic>Perceptual), and a preference for sentences over nonwords during passive reading (β=0.52, p=.009), but its response during sentence reading is substantially weaker than during semantic tasks on sentences (β=0.6, p<.001). Thus, the angular gyrus fROI appears to be more task-driven than the vATL fROIs.

## Discussion

What brain regions support semantic reasoning—the process of accessing, selecting, and combining conceptual representations? Using precision fMRI, we searched for brain regions that respond during semantic reasoning over both sentences and pictures. We discovered a set of regions that satisfy this amodal semantics criterion. Located in left frontal and temporo-parietal cortex and in the right cerebellum, these regions are selective for semantic tasks, showing a low response during the control perceptual tasks, passive sentence reading, and a demanding spatial working memory task. Critically, these semantic regions are distinct from the language system, contrary to the idea that language serves as a vehicle for abstract thought. They are also distinct from two other higher-order cognitive networks: the multiple demand and default mode networks. This novel type of neural selectivity challenges the field’s current understanding of large-scale functional organization of the human brain, emphasizing the unique requirements of semantic reasoning that may demand its own functionally specialized neural hardware.

To overcome the limitations of anatomy-based inference, we developed a functional localizer—an experimental paradigm that targets the cognitive process of interest and can identify the relevant areas reliably within individuals. This ‘localizer’ paradigm can then be used consistently across studies, along with additional paradigms designed to answer diverse research questions about the areas of interest. In this way, the localizer paradigms allow for straightforward combining of knowledge across studies and labs, as needed for a cumulative research enterprise.

Our semantic>perceptual localizer contrast generalizes across stimuli and tasks and consistently—across three experiments—identifies a set of brain regions that show a signature of amodal semantic processing: strong responses during a semantic task on either sentences or pictures, and weak responses to challenging perceptual tasks on the same stimuli. The broad anatomical locations of these areas align well with prior studies, including those that have looked for overlap between verbal and pictorial stimuli (e.g., Fairhall & Caramazza, 2013; Vandenberghe et al., 1996; Wurm & Caramazza, 2019) and those that have examined tasks that require retrieval of particular kinds of semantic information (see Jackson, 2021 for a meta-analysis). However, establishing whether those contrasts engage the same areas as the ones identified here will require within-participant comparisons, to be carried out in future studies.

### A new type of functional selectivity

Our newly discovered set of brain areas show strong selectivity for the semantic compared to the perceptual task (responding ∼3 times more strongly to the semantic conditions), respond only weakly during passive sentence comprehension, and show little response to a non-semantic working memory task. This response profile differs critically from those of the language network (which responds more strongly to sentences across tasks and more so than to pictures), the multiple demand network (which responds to all cognitively demanding tasks—in this case, semantic, perceptual, and working memory), and the default mode network (which shows below-baseline responses to cognitively demanding tasks, including the semantic task).

Why might semantic reasoning rely on its own neural machinery? First, it operates over concepts, which are not necessarily linguistic in nature—hence the dissociation between semantic and language brain regions. Indeed, conceptual reasoning likely predates language both evolutionarily (e.g., Thompson, 1995; Zentall et al., 2014) and developmentally (e.g., Kabdebon & Dehaene-Lambertz, 2019; Younger, 2010), indicating that at least some neural machinery for conceptual thought should be in place before the language network arises in the primate lineage or develops in human infants. Second, the demands of semantic reasoning are distinct from domain-general task demands, such as working memory, and formal reasoning, such as mathematical and logical reasoning—hence the dissociation between the neural basis of semantic reasoning and formal reasoning. In addition to manipulating abstract cognitive variables, semantic reasoning relies on the vast amount of conceptual knowledge about the world (Binder & Desai, 2011; Kumar, 2021; Markovits & Barrouillet, 2002). This includes our knowledge of the physical world and the social world, which have now been shown to rely on specialized brain systems (intuitive physics: Fischer et al., 2016; Kean et al, 2025; Paulun et al., 2025; Pramod et al., 2025) (social cognition: Adolphs, 1999, 2009; Deen et al., 2015; Saxe, 2006). The relationship between those domain-specific reasoning systems and the semantic reasoning regions remains to be discovered. One possibility is that the semantic regions serve as an intermediate stage between modality-specific (e.g., linguistic or visual) processing and reasoning, with the reasoning systems assembling individual concepts and ideas into more complex conceptual structures. Another possibility is that the semantic regions operate at the same level of the cognitive hierarchy, with the specialized reasoning regions additionally recruited when particular kinds of content need to be processed.

### Semantic control vs. semantic representation

One line of research (Badre et al., 2005; Badre & Wagner, 2002; Jackson, 2021; Jackson et al., 2021; Lambon Ralph et al., 2017) has implicated regions in the inferior frontal gyrus and the posterior temporal lobe in semantic control, defined as the process of selecting relevant conceptual associations and suppressing irrelevant ones (for example, selecting the auditory features of a piano in a context of someone playing a sonata, or its size and weight in a context where the piano needs to be moved). Those semantic control regions are similar in their general anatomical location to the regions we report here (see **Figure S7, B**), thus providing convergent evidence for these regions’ role in semantic cognition. Our results align with prior work that showed separability of semantic control from multiple demand and default mode regions (Chiou et al., 2023) but additionally highlight the distinction between these regions and the language network.

We chose not to use the term “semantic control” because semantic reasoning plausibly requires more than just control. Before the relevant meanings are selected, the space of possible meanings needs to be defined (through associative activation or otherwise). And after the relevant meanings are selected, they need to be combined and transformed in order to yield the answer relevant to current goals. Further, deep semantic cognition may be recruited in the absence of an explicit task, e.g. when a person is engaged in a stream of thought or listening to an interesting story. Indeed, the semantic regions from this study are topographically similar to a set of brain regions reported in Ryskina et al (2025), where participants were asked to simply think about the meaning of words, without a specific task in mind (see **Figure S7, C**). If those areas are the same as the semantic regions from this study, then specific semantic demands are indeed not required for their engagement. Whether the semantic regions we identify are implicated in semantic control or semantic reasoning more generally remains to be determined.

The researchers who advocate for the existence of semantic control brain regions draw a distinction between semantic control and semantic representation (Lambon Ralph et al., 2017). Semantic representation areas (specifically, the bilateral ventral ATL) are hypothesized to store concepts, and are therefore expected to respond to semantic content regardless of task demands. The most well-known proposal arguing for an amodal semantic representation hub in the ATL is the “hub-and-spokes” hypothesis (Patterson et al., 2007). Evidence for the hub component of this proposal comes from patients with semantic dementia—a subtype of primary progressive aphasia, a degenerative brain disease that affects different brain systems, including those relevant to linguistic and semantic processing (Gorno-Tempini et al., 2004; Mesulam et al., 2014). Patients with semantic dementia exhibit deterioration of semantic knowledge, showing gradual loss of details of concepts (Hodges & Patterson, 2007). Importantly, this loss is apparent for diverse features of concepts and no matter how the knowledge is probed (e.g., asking questions using language vs. categorizing pictures vs. asking the patients to draw an object from memory; Rogers et al, 2004). However, whether these areas store all conceptual knowledge remains debated: the most compelling evidence comes from concrete objects (e.g., the categories of animals, tools, food). Some have explicitly argued that conceptual representations of actions and events are stored and processed elsewhere (Mirman et al., 2017; Schwartz et al., 2011).

Our work provides evidence that ventral ATL supports semantic representation not only for single objects, but also for events: semantic fROIs in left and right ventral ATL respond during the semantic task (on both pictures and sentences) as well as during passive sentence reading (see Y. Zhang et al., 2024, for convergent evidence). That response profile, however, is distinct from the semantic regions that emerged in our whole-brain analysis: those respond only weakly to passive sentence reading, indicating that semantic reasoning exerts additional demands on these areas. Thus, our results are consistent with the view that ventral ATL areas and semantic regions in the frontal and temporo-parietal cortex are functionally distinct.

### Language understanding beyond the language network

We observed a dissociation between the language network and the semantic regions: the language areas respond much more strongly to the condition that requires semantic processing of sentences compared to the one that requires semantic processing of pictures (see also Ivanova et al., 2021; Sueoka et al., 2024), whereas the amodal semantic regions respond similarly strongly to the two semantic conditions and don’t respond to the language localizer contrast based on passive reading of sentences (vs. nonword sequences), which strongly engages the language areas. This dissociation suggests that these two sets of areas support distinct cognitive processes.

Although the language areas have been implicated in semantic composition, along with lexical retrieval and syntactic structure building (e.g., Shain, Kean, et al, 2024), some evidence suggests that the processing of linguistic meanings in these areas is quite shallow. For example, the language areas respond similarly strongly to plausible sentences and well-formed but non-sensical sentences, such as the famous *Colorless green ideas sleep furiously* example (Humphries et al., 2006; Kauf et al., 2024). Discriminating such stimuli requires reliance on world knowledge, which the language network may not have (full) access to. It is therefore possible that the language areas perform basic processing of the linguistic signals—recognizing words (and larger constructions) and figuring out how words go together—and then, if needed, pass this information to various downstream systems that can process the meaning more deeply (see Casto et al, 2025 a, for discussion).

We have here focused on amodal semantic regions, but several other networks that support high-level cognition can, too, receive information from the language system, including the two domain-general systems mentioned earlier (multiple demand and default mode networks), as well as domain-specific cognitive systems, such as the theory of mind network (DiNicola et al., 2020; Saxe, 2006; Saxe, Moran, et al., 2006) and the intuitive physics network (Fischer et al., 2016; Pramod et al., 2025). These systems support different kinds of reasoning and are all distinct from the language network (Braga et al., 2019; Diachek et al., 2020; Du et al., 2024; Fedorenko & Blank, 2020; Kean et al, 2025; Shain, Paunov, Chen et al, 2023). A key difference between the language network and these reasoning systems is that the language system is language-selective, but these other systems perform their computations on both linguistic and non-verbal inputs (e.g., pictures, digits, etc.; Casto et al, 2025 a; Fedorenko et al., 2024). Whether and how these higher-order cognitive networks interact with the semantic regions described here is an important area of future research.

### Cerebellar engagement

The semantic selectivity profile manifested itself not only in the left-hemisphere neocortical areas, but also in the right cerebellum. This evidence is in line with accounts of cerebellar involvement in cognitive function in general (e.g., Ito, 2008; Jacobi et al., 2021; Rapoport et al., 2000; Schmahmann, 2019; Schmahmann et al., 2019; Strick et al., 2009) and language in particular (Casto et al, 2025 b; Fedorenko et al., 2011; Highnam & Bleile, 2011; LeBel et al., 2021; Mariën & Borgatti, 2018; Smet et al., 2007; Starowicz-Filip et al., 2017), as well as reports of functional connectivity between cerebellar regions and neocortical networks, including multiple demand and default mode networks (Buckner et al., 2011b; Guell, Gabrieli, et al., 2018; Guell, Schmahmann, et al., 2018; for a review, see Habas, 2021). As in the neocortex, the response profile of the semantic cerebellar fROIs we described (Cer1 and Cer2) is clearly distinct from language-selective areas and domain-general multiple demand areas previously reported in the cerebellum (Casto et al, 2025 b; Guell, Gabrieli, et al., 2018; Schmahmann et al., 2019). Thus, our findings also contribute to the body of work aiming to discover the organization of the cognitive cerebellum.

### The importance of the left hemisphere for abstract semantic reasoning

A striking feature of the current results is the lateralization of the semantic regions to the left hemisphere (and to the right cerebellum, which is connected to the left neocortex). Not only were the semantic regions themselves left-lateralized, but to the extent that we saw a semantic>perceptual effect in other brain networks—multiple demand and default mode—it was only present in their left-hemisphere components. This left-hemispheric bias has plausibly led to the conflation of semantic and linguistic processing in some past studies, given that a left-hemispheric bias is often taken as evidence that the language system is engaged. However, many functions that are distinct from language show a left-hemispheric bias: from mathematical reasoning (Amalric & Dehaene, 2016; Istomina & Arsalidou, 2024) to logic (Kean et al, 2025; Monti et al., 2007, 2009) to intuitive physical reasoning (Kean et al, 2025; cf. Fischer et al., 2016). Apparently, abstract semantic reasoning is another such function.

What might be the reason for the left-hemispheric bias in semantic cognition? We learn about the world from two main sources: our lived experience (absorbing the world through our senses and physically interacting with the world) and language. In fact, most of the stuff you know, you have probably learned through language, without any direct experience. Amodal conceptual representations should be easily accessible both a) from perceptual modalities (which are bilaterally present) but also b) through language (which is left-lateralized), so the lateralization of language may well drive the location of these amodal semantic regions. Whether semantic regions reside in the right hemisphere in individuals with right-lateralized language systems will provide one test of this hypothesis.

### Implications for AI

The dissociation between linguistic and semantic processing has important implications for understanding the abilities and internal representations of large language models (LLMs). Although many LLMs are trained on language alone, their capabilities extend beyond language processing and today include various forms of reasoning (Huang & Chang, 2023; Plaat et al., 2025). The closest parallel to semantic reasoning in the LLM literature is the term “commonsense reasoning” (e.g., Davis & Marcus, 2015), which LLMs exhibit mixed success on. For example, small-to-medium LLMs are better at using social concepts (e.g., “friend” or “help”) compared to physical and spatial concepts (e.g., “left” or “inside”; Ivanova, Sathe, Lipkin et al, 2025) and can struggle to distinguish events based on real-world plausibility (Kauf, Ivanova et al, 2023; Leivada et al., 2025). Small language models trained on developmentally realistic amounts of data struggle with reasoning over semantic concepts in context, even though they successfully master linguistic rules (M. Hu et al., 2024). Thus, LLMs also show evidence of a dissociation between linguistic and conceptual semantic knowledge.

Semantic reasoning can be considered a type of functional linguistic competence—a non-language-specific skill that is nevertheless essential for competent language use in real-life settings (Mahowald, Ivanova et al, 2024). The current work emphasizes the non-language-specificity of semantic reasoning by explicitly imposing the requirement that the semantic regions should be sensitive to semantic content in both sentences and pictures. We expect that humanlike AI systems will exhibit a similar dissociation between linguistic and semantic processing (whether built-in or emerging organically over the course of training). In particular, the newly popular vision-language models (VLMs; J. Zhang et al., 2024) can be considered to mimic human processing if they develop language-specific and vision-specific processing steps, which then feed into a shared semantic processing stage.

### Open questions

The discovery of functionally selective semantic regions raises a host of important questions for further clarifying their function:

1. What counts as semantic reasoning? We show that two very different tasks—reversibility and plausibility—both recruit the semantic regions. What other tasks might or might not recruit them? Do these areas generalize to semantic content beyond actions (for a related meta-analysis, see Kuhnke, Beaupain, et al., 2023)? Are there instances when these regions contribute to task-free naturalistic cognition as well?
2. Do the semantic regions we identify form a single functionally cohesive network, like the language regions, or are they functionally heterogeneous, contributing to semantic cognition in distinct ways?
3. What is the representational format that these regions employ? Is there a shared representational format across input types and tasks or are do neural contain traces of their source modality and/or tasks? (see Fairhall & Caramazza, 2013; Wurm & Caramazza, 2019, for evidence that posterior temporal but not frontal areas contain generalizable crossmodal information).

The ability to sift through world knowledge to retrieve relevant information in a matter of seconds is a remarkable cognitive feat. We here have provided evidence that this complex process may require dedicated neural machinery, dissociable from the neural substrate of language processing. Our findings highlight the importance of precision brain imaging for uncovering functional dissociations in the human brain and pave the way for a neurocognitive exploration of the mechanisms underlying semantic reasoning.

## Method

### Participants

We collected data from 14 participants for Experiment 1 (11 women, 3 men, mean age = 25.9 years, SD = 8.10), 18 participants for Experiment 2 (7 women, 11 men, mean age = 21.0 years, SD = 1.68), and 26 participants for Experiment 3 (11 women, 13 men, mean age = 23.4 years, SD = 4.73). The participants were recruited from MIT and the surrounding Cambridge/Boston, MA, community and paid for their participation. All were native speakers of English, had normal or corrected to normal vision, and no history of language impairment. Two participants from Experiment 1, one from Experiment 2, and three from Experiment 3 were excluded due to low data quality; two participants from Experiment 2 and one participant from Experiment 3 were further excluded due to low behavioral task performance, leaving a total of 12, 15, and 22 participants for Experiments 1, 2, and 3, respectively. The protocol for the study was approved by MIT’s Committee on the Use of Humans as Experimental Subjects (COUHES). All participants gave written informed consent in accordance with protocol requirements.

A key distinguishing feature of our work compared to previous investigations of amodal semantics is the use of the individual-subject functional localization method (Fedorenko et al., 2010; Saxe, Brett, et al., 2006). This approach, falling within the broader methodology of precision fMRI, stands in contrast to traditional group-averaging analyses, whereby neural responses are averaged across participants on a voxel-by-voxel basis, and the resulting activation clusters are interpreted via ‘reverse inference’ from anatomy (e.g., Fedorenko, 2021; Poldrack, 2006, 2011). Group analyses tend to overestimate overlap in cases of nearby functionally distinct areas (Nieto-Castañón & Fedorenko, 2012), which is problematic for studies that aim to establish regions of shared activations across input modalities. This consideration is particularly important when analyzing responses in association cortex, where different functional regions vary in their precise locations across individuals (Blank et al., 2017; Fedorenko & Kanwisher, 2009; Frost & Goebel, 2012; Shashidhara, Spronkers, et al., 2019; Tahmasebi et al., 2012; Vázquez-Rodríguez et al., 2019) and often lay side by side in the form of ‘interdigitated networks’ (Braga et al., 2019, 2020; Deen & Freiwald, 2022; DiNicola et al., 2020; Fedorenko et al., 2012). The use of functional localization allows us to distinguish between semantic, linguistic, and domain-general executive processes even if they recruit brain regions that are adjacent to each other, thus enabling powerful and generalizable inferences about the neural basis of semantic reasoning.

### Critical experiments

The critical experiment in all three studies had a 2x2 blocked design: in each block, the stimuli were either sentences or pictures, and the task was either semantic (main) or perceptual (control). Thus, each experiment had four conditions: sentences + semantic task (SENT_SEM), sentences + perceptual task (SENT_PERC), pictures + semantic task (PIC_SEM), and pictures + perceptual task (PIC_PERC). To test the generalizability of our findings, we varied the nature of the stimuli and tasks across experiments. During behavioral piloting, the semantic and perceptual tasks within each experiment were adjusted to be approximately matched in difficulty.

#### Experiment 1 (*Figure 1A*, left)

The materials consisted of 120 pairs of color photographs depicting interactions of people with everyday objects (except one pair that depicted animate-animate interactions), as well as 120 pairs of ‘corresponding’ sentences that described the same interactions. Half of the stimuli depicted/described plausible scenarios (e.g., “peeling a carrot”), and half of the stimuli depicted/described implausible scenarios (e.g., “peeling a candle”). The photographs in each pair were highly similar except for the object used. Furthermore, materials were created in quadruplets, so that any given object was used in a plausible version in one pair, and in an implausible version in another pair (for example, for the peeling a candle / carrot, there was another pair: lighting up a carrot / candle).

The semantic task was to determine whether a given event was plausible or implausible. The perceptual task was to determine whether the stimulus was moving left or right (see below).

Before the experiment, participants completed a practice run outside the MRI scanner with 8 picture and 8 sentence stimuli (which were never shown inside the scanner). During the experiment, each of the 480 unique stimuli (240 pictures and 240 sentences) was presented twice: once in the semantic task, and once in the perceptual task, for a total of 960 trials. The 960 trials were split into 5 subsets of 192 trials each (48 for each of the four conditions), corresponding to five runs, with the constraint that any given event (e.g., a man peeling a carrot) occurred only once within a run (i.e., in only one of the four conditions). The 192 trials in each run were further grouped into 16 blocks (4 per condition) of 12 trials each, ensuring that the plausible and the implausible versions of the same event did not occur within the same block (within or across the four conditions). The exact grouping of the trials varied across participants.

Each trial lasted 2 s and consisted of the stimulus presented for 1.4 s followed by 0.6 s of fixation. During stimulus presentation (across all four conditions), the stimulus appeared on the screen (centered vertically, with the horizontal position drawn uniformly from an 80 pixel range around the vertical midline of the screen) and then moved diagonally toward the top left or top right corner. The picture and sentence stimuli moved at the velocity of (1,10) and (2,10) pixels in (x,y)-direction, respectively (the speed was chosen so as to approximately match the semantic and perceptual conditions for difficulty, based on behavioral piloting). Each block was preceded by a 2 s instructions screen to tell participants whether to perform the semantic or the perceptual task on the pictures/sentences that follow, and to remind them which button to press for which response: “1 – plausible; 2 – implausible” or “1 – left; 2 – right”. The instructions were additionally displayed in small font in the top right corner throughout each block.

Each experimental block lasted 26 s (2 s instructions + 12 trials * 2 s each). Each run consisted of 16 experimental blocks (4 per condition) and 5 fixation blocks each 16 s in duration, for a total duration of 496 s (8 min 16 s). A fixation block appeared at the beginning of each run, and after each set of four experimental blocks. Condition order was counterbalanced across runs and participants and was palindromic within each run (to counteract scanner drift). Each participant completed five runs for a total of 20 blocks per condition per subject. For consistency with the other two experiments, each of which only included two runs, we here focus on the first two runs for each participant (the responses for all five runs were attenuated due to adaptation but showed a similar pattern).

#### Experiment 2 (*Figure 1A*, center)

The materials consisted of color photographs and sentences depicting/describing interactions of people with everyday objects (except two photos and two sentences that depicted/described animate-animate interactions). Unlike in Experiment 1, the scenarios in the pictures and sentences were different (to minimize the possibility of cross-coding: i.e., recalling the verbal representation for a picture if a sentence version of the same event has been previously encountered, or recalling the pictorial representation for a sentence if a picture version has been previously encountered). Further, all scenarios were plausible (to ensure that responses in Experiment 1 are not driven by the unexpected nature of half of the stimuli), but some depicted/described irreversible actions, i.e., actions, the effects of which cannot be undone (e.g., eating a clementine or peeling a carrot) and others depicted/described reversible actions (e.g., putting staples into a stapler or inserting a key into a lock). A subset of the plausible photographs from Experiment 1 (N=112) were reused for this study along with newly created sentences (N=112).

The semantic task was to determine whether a given action was reversible or irreversible. The perceptual task was to determine whether the movement direction of the stimulus changed 3 or 4 times (two participants completed an earlier version of the task where the direction changed either 5 or 7 times).

Before the experiment, participants completed a practice run outside the scanner with 8 picture and 8 sentence stimuli (not used inside the scanner). For the main experiment, we created two lists, where each stimulus was assigned either to the semantic condition or to the perceptual condition. Across the two lists, we balanced the ratio of actions irreversibly affecting the object to those reversibly doing so and the split of these event types across modalities (i.e., each list consisted of 83 irreversible events, and 141 reversible events; 43 of the 83 irreversible events were presented as pictures and 40 were presented as sentences). Each participant was assigned one of the two lists. Each list was split into two subsets of 112 trials each (28 for each of the four conditions), corresponding to two runs. The 112 trials in each run were further grouped into 16 blocks (4 per condition) of 7 trials each, ensuring that each block contained between 1 and 4 irreversible events and that at most 3 trials in a row changed direction the same number of times.

Each trial lasted 2 s and consisted of the stimulus presented for 1.4 s followed by a prompt (“RESPOND”) presented in the center of the screen in red font for 0.6 s. During stimulus presentation (across all four conditions), the stimulus appeared in the center of the screen and then randomly changed directions. The picture and sentence stimuli moved at the velocity of (1,10) and (2,10) pixels in (x,y)-direction, respectively (the speed was chosen so as to approximately match the semantic and perceptual conditions for difficulty, based on behavioral piloting). Each block was preceded by a 2 s instructions screen to tell participants whether to perform the semantic or the perceptual task on the pictures/sentences that follow, and to remind them which button to press for which response: “1 - irreversible; 2 – reversible” or “1 – odd number of direction changes; 2 – even number of direction changes”. The instructions were additionally displayed in small font in the bottom left corner throughout each block.

Each experimental block lasted 16 s (2 s instructions + 7 trials * 2 s each). Each run consisted of 16 experimental blocks (4 per condition) and 5 fixation blocks each 16 s in duration, for a total duration of 336 s (5 min 36 s). A fixation block appeared at the beginning of each run, and after each set of four experimental blocks. As in Experiment 1, condition order was counterbalanced across runs and participants, and was palindromic within each run. Each participant completed two runs.

#### Experiment 3 (*Figure 1A*, right)

This experiment was previously reported in Ivanova et al (2021).The materials consisted of 40 pairs of line drawings depicting interactions between two animate entities, as well as 40 pairs of ‘corresponding’ sentences that described the same interactions. As in Experiment 1, half of the stimuli depicted/described plausible scenarios (e.g., “a cop is arresting a criminal”), and half of the stimuli depicted/described implausible scenarios (e.g., “a criminal is arresting a cop”). The implausible events were created by swapping the agent and the patient of the plausible events. The semantic task was to determine whether a given event was plausible or implausible, and the perceptual task was to determine whether the stimulus was moving left or right (as in Experiment 1).

As in Experiment 2, we created two lists, where each stimulus was assigned either to the semantic condition or to the perceptual condition. Each list was split into two subsets of 80 trials each (20 for each of the four conditions), corresponding to two runs, ensuring that the same event (in picture vs. sentence format) does not occur in the same run. The 80 trials in each run were further grouped into 8 blocks (2 per condition) of 10 trials, ensuring that at most 3 plausible or implausible events appeared in a row. Movement direction of the stimuli was assigned randomly.

Each trial lasted 2 s and consisted of the stimulus presented for 1.5 s followed by 0.5 s of fixation. During stimulus presentation (across all four conditions), the stimulus appeared in the center of the screen and then moved horizontally toward the left or right. The picture and sentence stimuli both moved at the velocity of 0.025 pixels/screen in positive or negative x-direction (the speed was chosen to approximately match the semantic and perceptual conditions for difficulty, based on behavioral piloting). Each block was preceded by a 2 s instructions screen to tell participants whether to perform the semantic or the perceptual task on the pictures/sentences that follow, and to remind them which button to press for which response: “1 - plausible; 2 – implausible” or “1 - left; 2 – right”. The instructions were additionally displayed in small font in the bottom left corner throughout each block.

Each experimental block lasted 22 s (2 s instructions + 10 trials * 2 s each). Each run consisted of 8 experimental blocks (2 per condition) and 3 fixation blocks each 22 s in duration, for a total duration of 242 s (4 min 2 s). A fixation block appeared at the beginning of each run, and after each set of four experimental blocks. Condition order was counterbalanced across runs and participants and was palindromic within each run. Each participant completed two runs.

### Localizer experiments

In addition to the critical experiment, all participants completed the language localizer experiment (sentence and nonword list reading), designed to identify language-responsive regions in individual participants (Fedorenko et al., 2010). Participants in Studies 2 and 3 also completed a multiple demand network localizer experiment (a spatial working memory task), designed to identify the multiple demand regions in individual participants (Fedorenko et al., 2013). Data from the multiple demand network localizer was also used to define the default mode network (with a reverse contrast; see below).

#### Language network localizer

To localize the language network, we asked participants to read sentences (e.g., NOBODY COULD HAVE PREDICTED THE EARTHQUAKE IN THIS PART OF THE COUNTRY) and lists of unconnected, pronounceable nonwords (e.g., U BIZBY ACWORRILY MIDARAL MAPE LAS POME U TRINT WEPS WIBRON PUZ) in a blocked design. Each stimulus consisted of twelve words/nonwords. For details of how the language materials were constructed, see Fedorenko et al. (2010). The materials are available at https://evlab.mit.edu/funcloc/. The sentences > nonword-lists contrast isolates processes related to language comprehension (responses evoked by, e.g., visual perception and reading are subtracted out) and has been previously shown to reliably activate left-lateralized fronto-temporal language processing regions, be robust to changes in task and materials, and activate the same regions regardless of whether the materials were presented visually or auditorily (Fedorenko et al., 2010; Mahowald & Fedorenko, 2016; Scott et al., 2017). Further, a similar network emerges from task-free resting-state data (Braga et al., 2020). Stimuli were presented in the center of the screen, one word/nonword at a time, at the rate of 450 ms per word/nonword. Each stimulus was preceded by a 100 ms blank screen and followed by a 400 ms screen showing a picture of a finger pressing a button, and a blank screen for another 100 ms, for a total trial duration of 6 s. Participants were asked to press a button whenever they saw the picture of a finger pressing a button. This task was included to help participants stay alert. Condition order was counterbalanced across runs. Experimental blocks lasted 18 s (with 3 trials per block), and fixation blocks lasted 14 s. Each run (consisting of 5 fixation blocks and 16 experimental blocks) lasted 358 s. Each participant completed 2 runs.

#### Multiple demand and default mode network localizer

To identify the multiple demand system within individual participants, we asked them to complete a spatial working memory task with two difficulty conditions. Participants had to keep track of four (easy condition) or eight (hard condition) sequentially presented locations in a 3 × 4 grid (**Figure 1B**; Fedorenko et al., 2011). In both conditions, they performed a two-alternative forced-choice task at the end of each trial to indicate the set of locations they just saw. The hard > easy contrast has been previously shown to reliably activate bilateral frontal and parietal multiple demand regions (Assem et al., 2020; Blank et al., 2014; Fedorenko et al., 2013). Numerous studies have shown that the same brain regions are activated by diverse executively-demanding tasks (Duncan & Owen, 2000; Fedorenko et al., 2013; Hugdahl et al., 2015; Shashidhara, Mitchell, et al., 2019; Woolgar et al., 2011). The easy > hard contrast has been shown to localize default mode regions (Blank & Fedorenko, 2020; Mineroff et al., 2018), which are generally attenuated in the presence of an external task (e.g., Buckner et al., 2008; Buckner & DiNicola, 2019; Fox et al., 2005; Raichle, 2015).

Stimuli were presented in the center of the screen across four steps. Each step lasted 1 s and revealed one location on the grid in the easy condition, and two locations in the hard condition. Each stimulus was followed by a choice-selection step, which showed two grids side by side. One grid contained the locations shown across the previous four steps, while the other contained an incorrect set of locations. Participants were asked to press one of two buttons to choose the grid that showed the correct locations. Condition order was counterbalanced across runs. Experimental blocks lasted 32 s (with 4 trials per block), and fixation blocks lasted 16 s. Each run (consisting of 4 fixation blocks and 12 experimental blocks) lasted 448 s. Each participant completed 2 runs.

### fMRI data acquisition

Structural and functional data were collected on the whole-body, 3 Tesla, Siemens Trio scanner with a 32-channel head coil, at the Athinoula A. Martinos Imaging Center at the McGovern Institute for Brain Research at MIT. T1-weighted structural images were collected in 176 sagittal slices with 1mm isotropic voxels (TR=2,530ms, TE=3.48ms). Functional, blood oxygenation level dependent (BOLD), data were acquired using an EPI sequence (with a 90° flip angle and using GRAPPA with an acceleration factor of 2), with the following acquisition parameters: thirty-one 4mm thick near-axial slices acquired in the interleaved order (with 10% distance factor), 2.1mm×2.1mm in-plane resolution, FoV in the phase encoding (A>>P) direction 200mm and matrix size 96mm×96mm, TR=2000ms and TE=30ms. The first 10s of each run were excluded to allow for steady state magnetization.

### fMRI data preprocessing

fMRI data were analyzed using SPM12 (release 7487), CONN EvLab module (release 19b), and other custom MATLAB scripts. Each participant’s functional and structural data were converted from DICOM to NIFTI format. All functional scans were coregistered and resampled using B-spline interpolation to the first scan of the first session (Friston et al., 1995). Potential outlier scans were identified from the resulting subject-motion estimates, as well as from BOLD signal indicators, using default thresholds in CONN preprocessing pipeline (5 standard deviations above the mean in global BOLD signal change, or framewise displacement values above 0.9 mm; Nieto-Castañon, 2020). Functional and structural data were independently normalized into a common space (the Montreal Neurological Institute [MNI] template; IXI549Space) using SPM12 unified segmentation and normalization procedure (Ashburner & Friston, 2005) with a reference functional image computed as the mean functional data after realignment across all timepoints omitting outlier scans. The output data were resampled to a common bounding box between MNI-space coordinates (−90, −126, −72) and (90, 90, 108), using 2mm isotropic voxels and 4th order spline interpolation for the functional data, and 1mm isotropic voxels and trilinear interpolation for the structural data. Last, the functional data were smoothed spatially using spatial convolution with a 4 mm FWHM Gaussian kernel.

### First-level analysis

Responses in individual voxels were estimated using a General Linear Model (GLM) in which each experimental condition was modeled with a boxcar function convolved with the canonical hemodynamic response function (HRF) (fixation was modeled implicitly, such that all timepoints that did not correspond to one of the conditions were assumed to correspond to a fixation period). Temporal autocorrelations in the BOLD signal timeseries were accounted for by a combination of high-pass filtering with a 128 seconds cutoff and whitening using an AR(0.2) model (first-order autoregressive model linearized around the coefficient a=0.2) to approximate the observed covariance of the functional data in the context of Restricted Maximum Likelihood estimation (ReML). In addition to experimental condition effects, the GLM design included first-order temporal derivatives for each condition (included to model variability in the HRF delays), as well as nuisance regressors to control for the effect of slow linear drifts, subject-motion parameters, and potential outlier scans on the BOLD signal.

### Defining functional regions of interest (fROIs)

The critical analyses were restricted to individually defined functional regions of interest (fROIs). These fROIs were defined using the group-constrained subject-specific (GcSS) approach (Fedorenko et al., 2010; Julian et al., 2012), where a set of ‘parcels’ (masks) is combined with each individual subject’s activation map for the relevant contrast to constrain the definition of individual fROIs.

The parcels delineate the expected gross locations of activations for a given contrast and are sufficiently large to encompass the extent of variability in the locations of individual activations. We used two types of parcels: (a) semantic parcels defined based on the critical experiments from this study, and (b) three sets of parcels for well-established and characterized large-scale networks defined based on fMRI activation maps from the relevant localizer experiments: the language network, the multiple demand network, and the default mode network (DMN), all of which might contribute to semantic processing, as discussed in the Introduction. The semantic parcels were generated based on the data from 30 participants from this experiment (10 from each critical experiment, in order to provide a balanced representation of the three experiments). We used a subset of the data to be able to test whether our results generalize to left-out participants. The parcels were generated using a conjunction contrast of SENT_SEM > SENT_PERC (a contrast that targets semantic processing in sentences) and PIC_SEM > PIC_PERC (a contrast that targets semantic processing in pictures). The language network parcels were generated from previously collected language localizer data from 220 participants (using the sentences>nonwords contrast). The multiple demand and default mode network parcels were from previously collected spatial working memory task data from 197 participants (using the hard>easy and easy>hard contrasts, respectively). For details of the parcel definition procedure, see Fedorenko et al (2010).

Within each parcel, we selected the top 10% most responsive voxels, based on the *p-*values for the contrast(s) of interest. This top n% approach ensures that the fROIs can be defined in every participant, thus enabling us to generalize the results to the entire population (Nieto-Castañón & Fedorenko, 2012). The contrasts used to define the fROIs were the same as those used to create the parcels. For a simple contrast, we ranked the voxels based on the p-value for that contrast; for a conjunction contrast (i.e., SENT_SEM > SENT_PERC and PIC_SEM > PIC_PERC), we ranked the voxels based on the larger of the two *p*-values (which is equivalent to a soft ‘and’ conjunction and allows us to specify the exact number of voxels to be selected).

### Examining the functional response profiles of fROIs

After defining fROIs in individual participants, we measured their responses to conditions of interest by averaging the responses to each condition across voxels to get a single value per condition per fROI in each participant. The responses to the localizer conditions (e.g., sentences and nonwords for language fROIs) were estimated using an across-runs cross-validation procedure, where one run was used to define the fROI and the other to estimate the response magnitudes, then the procedure was repeated switching which run was used for fROI definition vs. response estimation, and finally the estimates were averaged to derive a single value per condition per fROI per participant. This cross-validation procedure allows one to use all of the data for defining the fROIs as well as for estimating their responses (see Nieto-Castañón & Fedorenko, 2012, for discussion), while ensuring the independence of the data used for fROI definition and response estimation (Kriegeskorte et al., 2009). One participant completed only one run of the multiple demand localizer task; therefore, we did not estimate the strength of their responses to the hard and easy multiple demand localizer conditions.

To compare responses across conditions, we ran linear mixed-effect regression models with participant as a random intercept (for analysis of responses at the network level, we also included fROI as a random intercept). The analysis was run using the *lmer* function from the *lme4* R package (Bates et al., 2015); statistical significance of the effects was evaluated using the *lmerTest* package (Kuznetsova et al., 2017). For most analyses, we jointly analyzed the data from the three critical experiments while adding *Experiment* and *Experiment*Condition* terms to the regression model to account for potential inter-experiment variability. All follow-up analyses used FDR correction for multiple comparisons; *p*-values were adjusted based on the number of groupings used in each case (e.g., number of fROIs for within-network fROI analyses, number of fROIs times the number of experiments for evaluating response consistency across experiments, two for within-network hemisphere comparisons).

### Overlap analysis

To determine whether our newly defined semantic fROIs overlap with fROIs from previously established networks, we calculated the overlap coefficient between each fROI pair (using the formula: number of voxels shared / number of voxels in the smaller fROI). As a control, we also calculated the overlap between fROIs defined using the same contrast, the same experiment, and the same parcel, but using data from two different runs.

## Acknowledgements

We thank Zachary Mineroff and Zuzanna Balewski for assistance with experimental design and data collection, Jin Li for feedback on this work, and members of Kanwisher and Fedorenko labs for their assistance throughout this project. NK and EF were supported by research funds from the McGovern Institute for Brain Research (including from the Poitras Center for Psychiatric Disorders Research), from the Siegel Family Quest for Intelligence, and from the Simons Center for the Social Brain.

## Data and code availability

The data required to reproduce the figures and the statistical analysis, as well as associated analysis code are available on Github: https://github.com/neuranna/TripleEvents. The semantic parcels are available on the LIT lab website: https://www.language-intelligence-thought.net/resources.

## Supplemental Information

### SI-1. Behavioral results

The behavioral results for the three critical experiments are shown in **Figure S1**. In the figure and in the text below, SENT = sentence stimuli, PIC = picture stimuli, SEM = semantic task, PERC = perceptual task.

#### Experiment 1

Average response rate was 0.80. Overall mean reaction time was 1.02s, SD=0.22 (SENT_SEM: 1.09s, SD=0.24; SENT_PERC: 1.01s, SD=0.23, PIC_SEM: 0.98s, SD=0.18, PIC_PERC: 1.01s, SD=0.21). RTs were slightly higher for the semantic task than for the perceptual task (β=0.02, SE=0.01, p<.001) and for sentence stimuli compared to picture stimuli (β=0.02, SE=0.01, p<.001), with an interaction between stimulus type and task (β=0.1, SE=0.01, p<.001). Overall mean accuracy was 0.82, SD=0.39 (SENT_SEM: 0.86, SD=0.35; SENT_PERC: 0.73, SD=0.45, PIC_SEM: 0.89, SD=0.31, PIC_PERC: 0.79, SD=0.41). Accuracy was higher for the semantic task than for the perceptual task (β=0.88, SE=0.09, p<.001) and lower for sentence stimuli compared to picture stimuli (β=-0.28, SE=0.09, p<.001), with no interaction between stimulus type and task.

#### Experiment 2

Average response rate was 0.89. Overall mean reaction time was 1.51s, SD=0.49 (SENT_SEM: 1.41s, SD=0.56; SENT_PERC: 1.62s, SD=0.44, PIC_SEM: 1.43s, SD=0.42, PIC_PERC: 1.57s, SD=0.49). RTs for the semantic task were slightly lower than for the perceptual task (β=-0.17, SE=0.02, p<.001), with an interaction between stimulus type and task (β=-0.08, SE=0.03, p=0.016). There was no main effect of stimulus type on reaction time.

Overall mean accuracy was 0.77, SD=0.42 (SENT_SEM: 0.64, SD=0.48; SENT_PERC: 0.84, SD=0.36, PIC_SEM: 0.77, SD=0.42, PIC_PERC: 0.84, SD=0.37). Accuracy was lower for the semantic task than for the perceptual task (β=-0.79, SE=0.09, p<.001) and for sentence stimuli compared to picture stimuli (β=-0.33, SE=0.09, p<.001), with an interaction between stimulus type and task (β=-0.75, SE=0.19, p<.001).

#### Experiment 3

Average response rate was 0.90. Overall mean reaction time was 1.09s, SD=0.52 (SENT_SEM: 1.12s, SD=0.6; SENT_PERC: 1.03s, SD=0.51, PIC_SEM: 1.14s, SD=0.47, PIC_PERC: 1.06s, SD=0.5). RTs for the semantic task were slightly higher than for the perceptual task (β=0.09, SE=0.02, p<.001). There was no significant effect of stimulus type (sentences vs. pictures) and no interaction between stimulus type and task. Overall mean accuracy was 0.68, SD=0.47 (SENT_SEM: 0.81, SD=0.39; SENT_PERC: 0.78, SD=0.42, PIC_SEM: 0.75, SD=0.43, PIC_PERC: 0.73, SD=0.44). Accuracy was higher for the semantic task than for the perceptual task (β=0.16, SE=0.07, p=0.031) There was no significant effect of stimulus type and no interaction between stimulus type and task. For 16 participants, accuracy data from one run was erased because of a bug in the script.

Overall, there is no consistent trend in difficulty patterns across experiments, suggesting that the observed neural response patterns cannot be explained by between-condition differences in difficulty.

**Figure S1.**
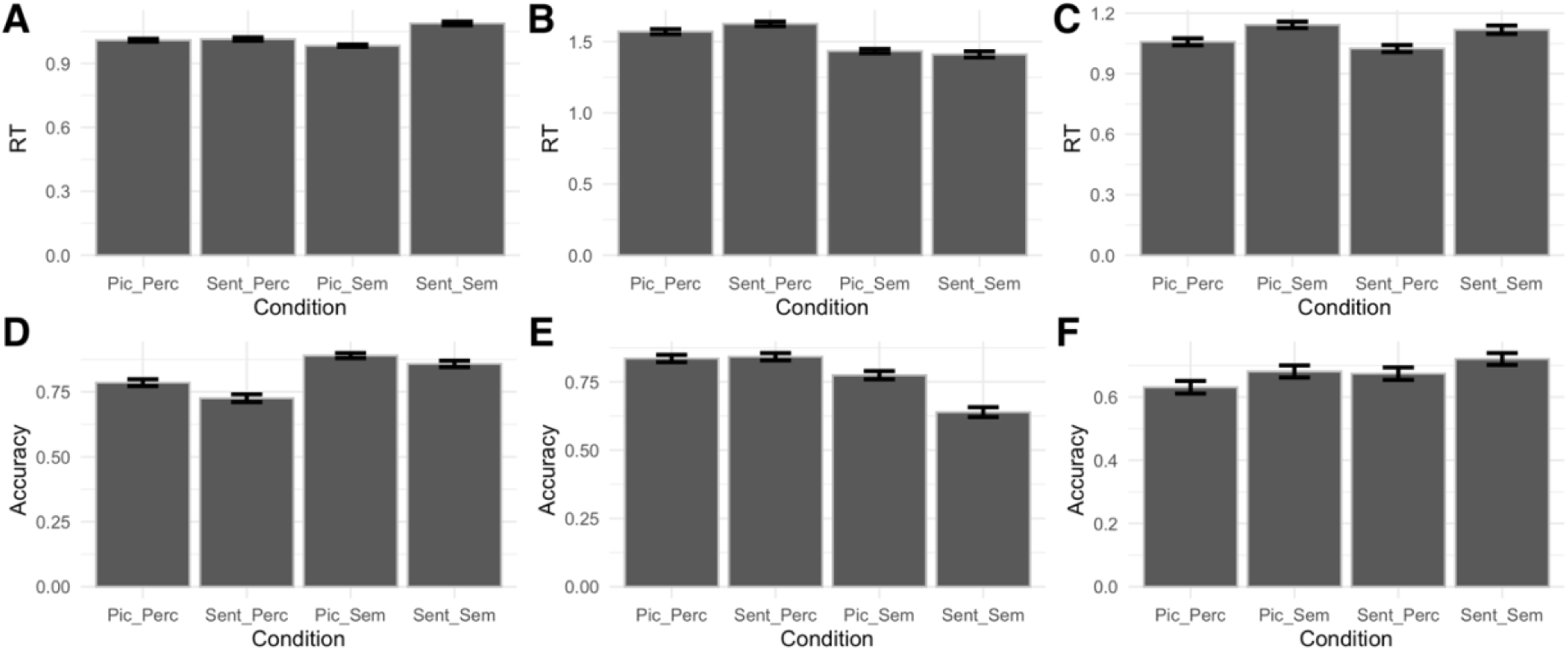
Behavioral results for the critical experiments. **(A-C)** Response times for Experiments 1, 2 and 3 respectively. **(D-F)** Accuracies for Experiments 1, 2, and 3, respectively.

### SI-2. Additional information about the responses in semantic fROIs

#### Semantic fROI responses generalize across experiments

Despite some variability in response magnitudes, overall response patterns are similar across experiments and fROIs (**Figure S2;** see main text for statistics).

**Figure S2.**
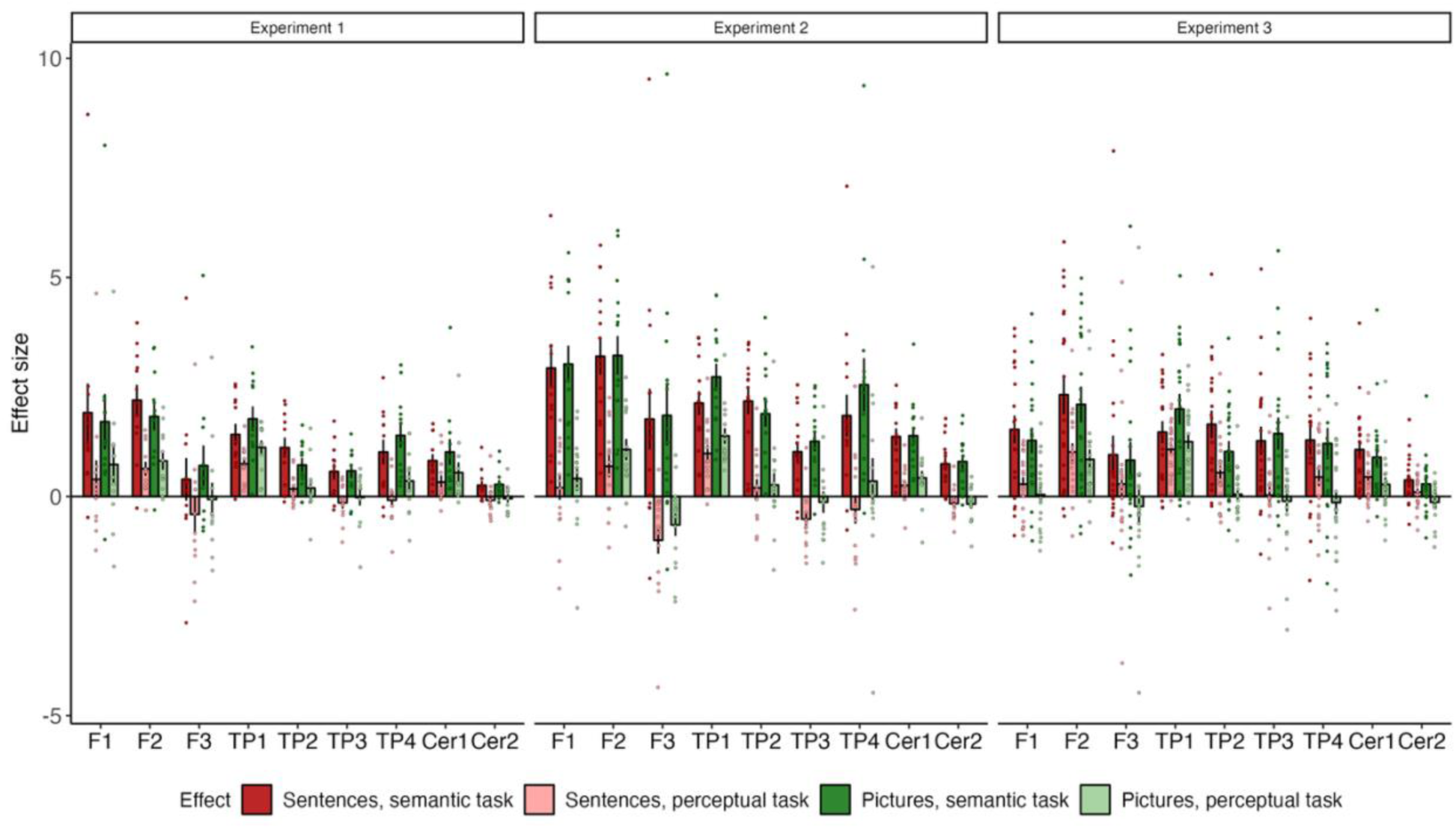
Semantic fROI responses in each experiment.

#### Visual parcels

Our whole-brain GcSS analysis yielded 11 parcels, 9 of which are described in the main text. Two parcels appear to be driven by low-level visual properties of the stimuli and are therefore excluded from the main analyses. Their locations and response profiles are shown in **Figure S3**.

**Figure S3.**
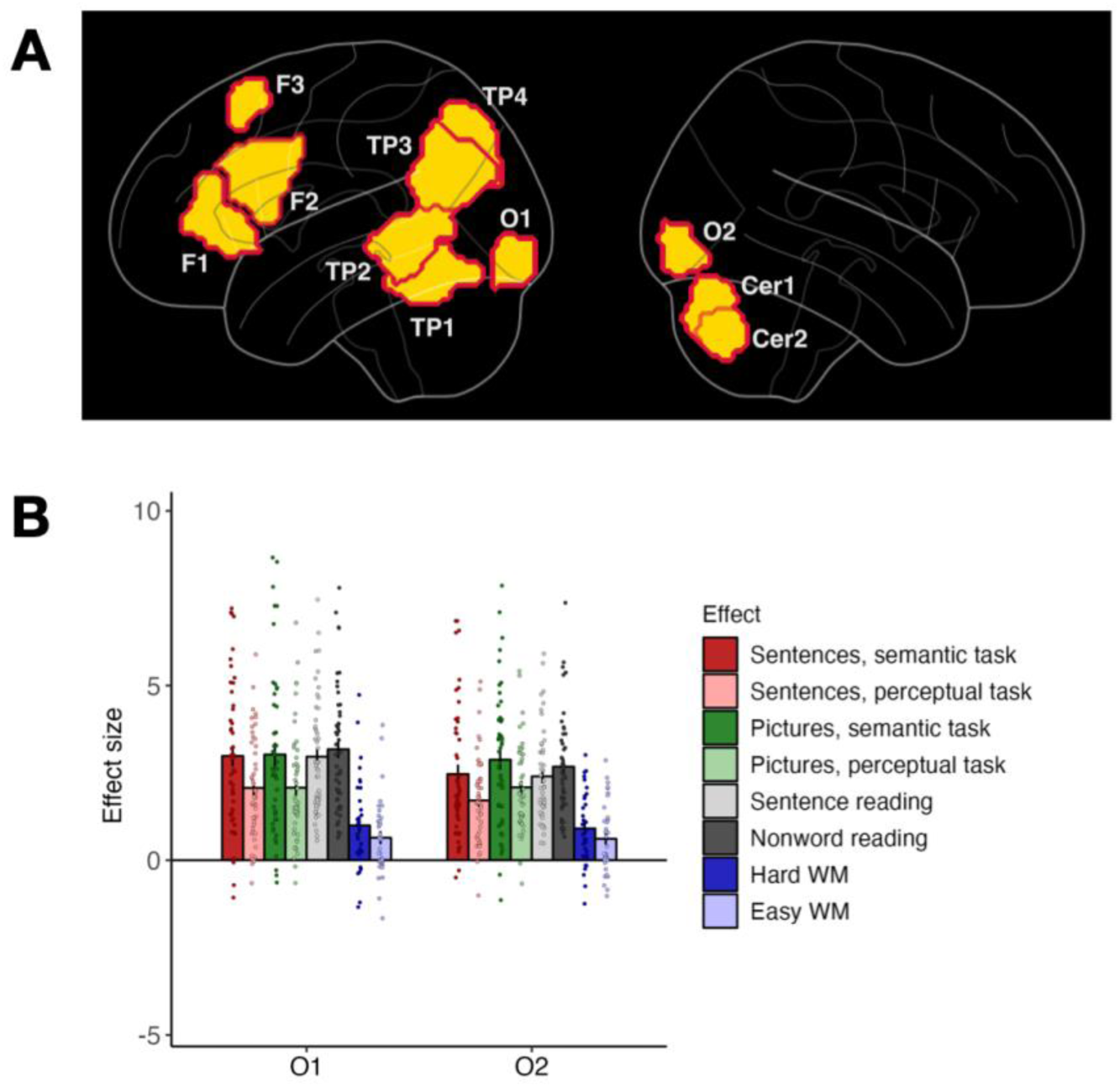
fROIs in bilateral early visual cortex were found as part of the brain-whole analysis but excluded from follow-up analyses due to our focus on cognitive rather than perceptual processing. **(A)** All parcels found via GcSS, including two in the occipital cortex: O1 (left hemisphere) and O2 (right hemisphere). **(B)** Response profiles of fROIs defined within the occipital parcels.

#### Semantic regions’ responses generalize to left-out participants

We use independent sets of data (different runs of the same task) when we define a fROI using data from a given experiment and measure its responses to the conditions from the same experiment. In addition, to test how well our results generalize to a new set of participants, we split the participants into two groups: the first group (n=30, 10 from each experiment) were used to define the parcels, and the second group (n=11) was left out. The results were consistent across the two groups: we observed a preference for semantic over perceptual tasks (both groups: p<.001), no overall effect of stimulus type, no interaction between stimulus type and task, no preference for sentences over nonwords or for hard vs. easy versions of the working memory task (**Figure S4**).

**Figure S4.**
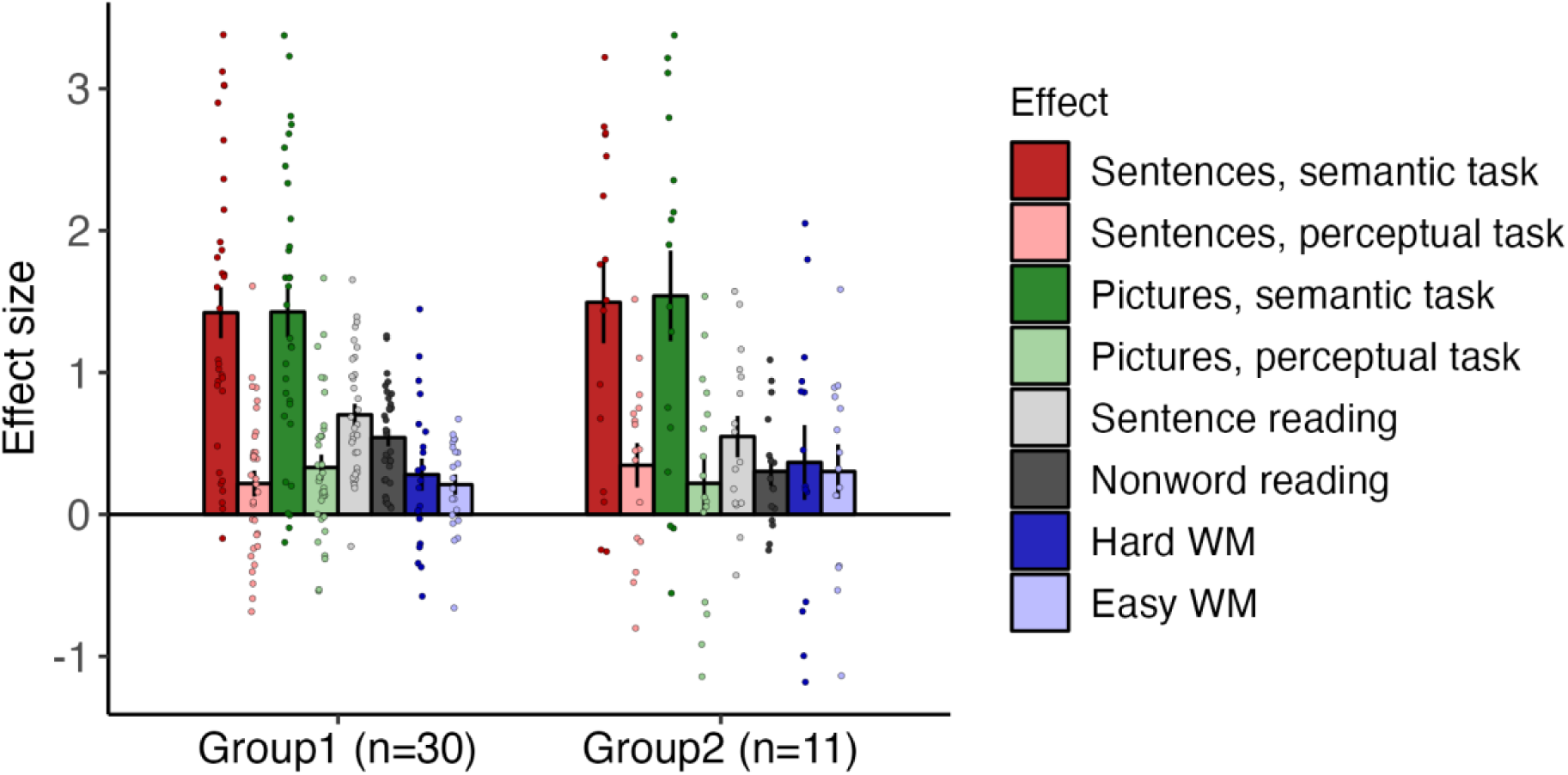
The results generalize to a new group of participants (whose data was not used for parcel definition). WM = working memory.

#### Contralateral semantic fROIs

We additionally examined responses in semantic parcels mirrored onto the right cerebral hemisphere and left cerebellum (the original parcels we found were located in the left cerebral hemisphere and right cerebellum). The response profiles of fROIs defined with these contralateral parcels are shown in **Figure S5**; see main text for summary and stats.

**Figure S5.**
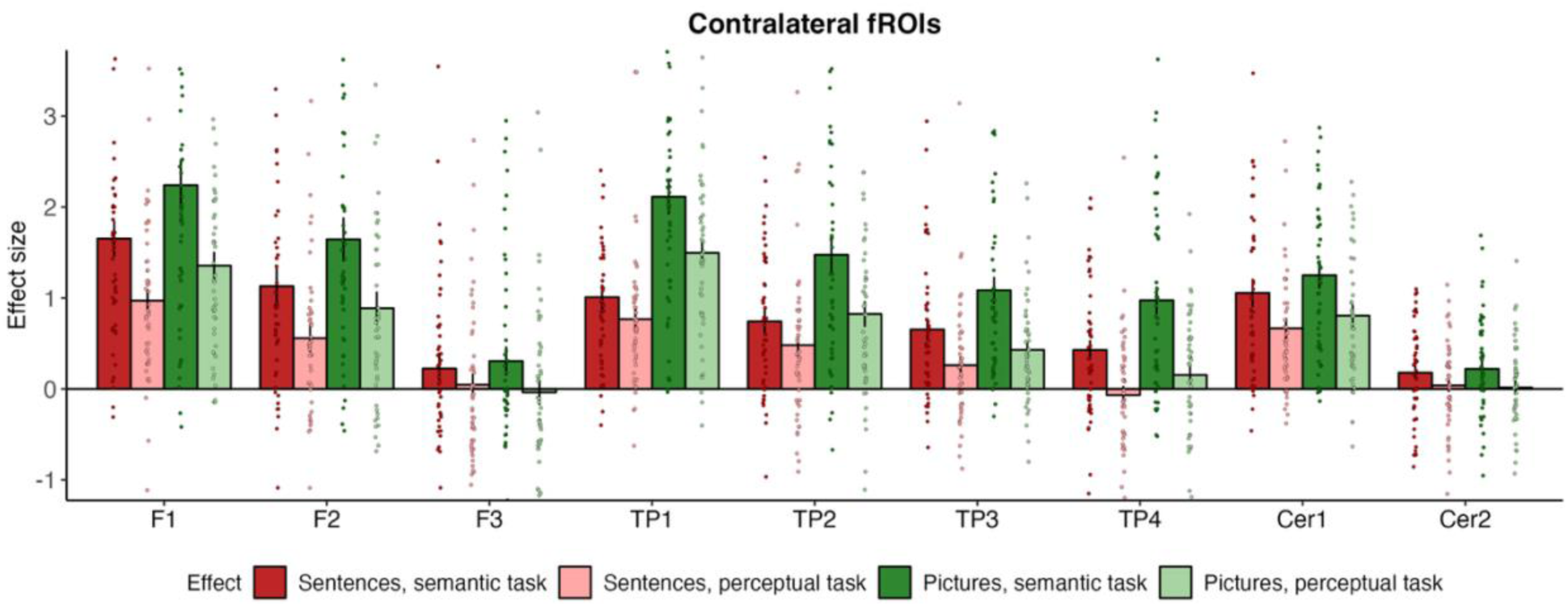
Response profiles of semantic fROIs defined using contralateral parcels (original semantic parcels projected onto the opposite hemisphere).

### SI-4. tSNR analysis details

To assess the temporal stability of the fMRI data, we calculated the temporal signal-to-noise ratio (tSNR) for each voxel across all subjects and functional runs (**Figure S6, A**). For each run, we used preprocessed data that underwent motion correction, spatial smoothing, and normalization to MNI space. We then generated voxel-wise tSNR maps by dividing the mean signal intensity by the temporal standard deviation of the signal at each voxel (Welvaert and Rosseel, 2013) and log transformed them for visualization purposes. To avoid division by zero, voxels with zero standard deviation were assigned a tSNR value of zero. A small constant (1e-10) was added to all tSNR values before log transformation to prevent undefined values.

Next, we calculated parcel-wise tSNR values to examine differences between semantic demand parcels and anatomical anterior temporal lobe (ATL) parcels. Semantic demand parcels were generated using the GcSS approach, while anatomical ATL parcels were adapted based on the Desikan Atlas (Desikan et al., 2006). Specifically, we selected the inf+pole+mid temporal lobe parcels and halved those anatomical regions along the anterior-posterior axis (y-axis) to isolate the anterior portion (**Figure S6, B**).

Using these anatomically defined parcels, we restricted the whole-brain tSNR maps to calculate the mean tSNR within each parcel. This allowed for a direct comparison of tSNR across regions of interest (**Figure S6, C**).

**Figure S6.**
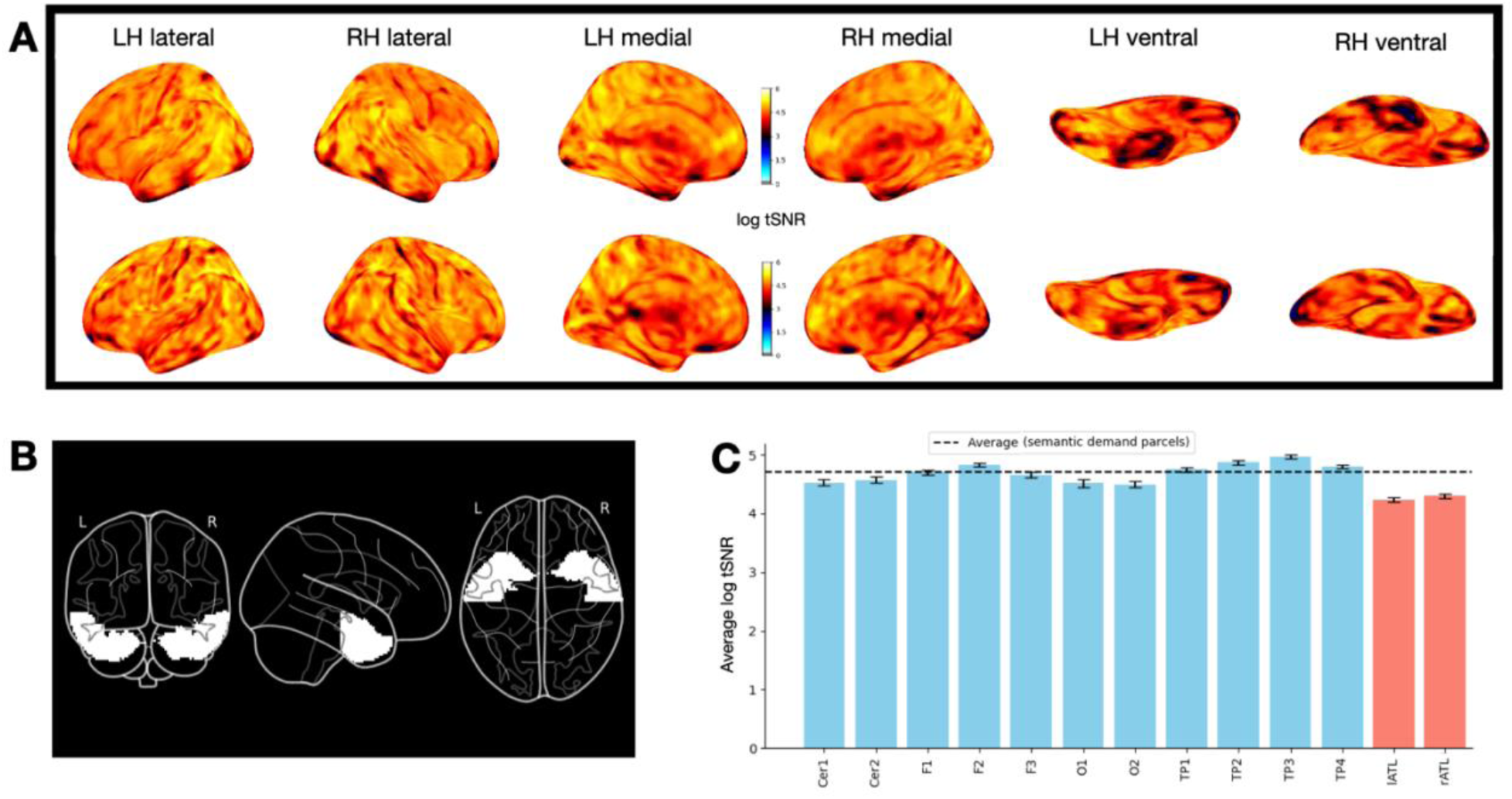
tSNR analysis. (**A**) Sample subject tSNR maps. (**B**) Anatomical ATL parcels. (**C**) Average tSNR in the semantic demand parcels discovered with whole-brain GSS analysis (blue) and in anatomically defined ATL parcels (red).

### SI-5. Comparison with previously described semantics-responsive brain regions

**Figure S7.**
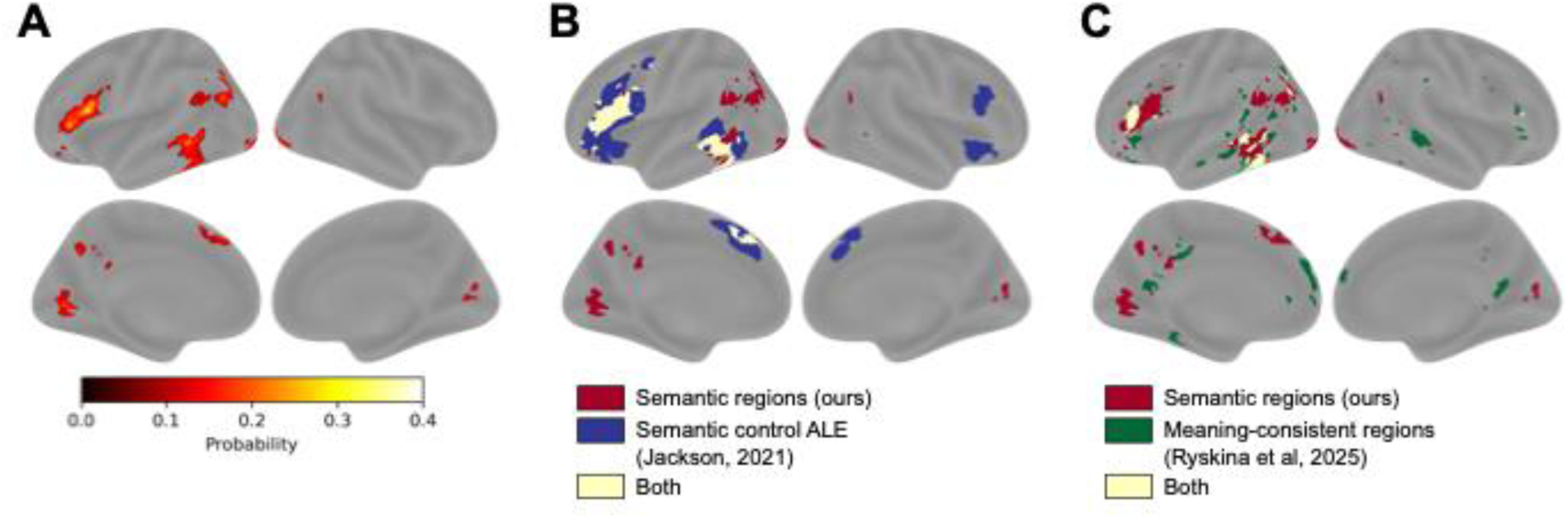
Group-level comparisons of semantic regions’ locations. **(A)** Probabilistic overlap map from which the semantic parcels were derived. Colorbar: ratio of participants who showed significant activation at this voxel. **(B)** Overlap between semantic regions (ours) and the semantic control regions derived from the meta-analysis in Jackson (2021). ALE = activation likelihood estimation. **(C)** Overlap between semantic regions (ours) and the meaning-consistent regions from Ryskina et al (2025).

