## Supplemental Info for "Semantic reasoning takes place largely outside the language network"

### Supplemental Information

#### SI-1. Behavioral results

The behavioral results for the three critical experiments are shown in **Figure S1**. In the figure and in the text below, SENT = sentence stimuli, PIC = picture stimuli, SEM = semantic task, PERC = perceptual task.

*Experiment 1.* Average response rate was 0.80. Overall mean reaction time was 1.02s, SD=0.22 (SENT_SEM: 1.09s, SD=0.24; SENT_PERC: 1.01s, SD=0.23, PIC_SEM: 0.98s, SD=0.18, PIC_PERC: 1.01s, SD=0.21). RTs were slightly higher for the semantic task than for the perceptual task ($\beta$=0.02, SE=0.01, $p<.001$) and for sentence stimuli compared to picture stimuli ($\beta$=0.02, SE=0.01, $p<.001$), with an interaction between stimulus type and task ($\beta$=0.1, SE=0.01, $p<.001$). Overall mean accuracy was 0.82, SD=0.39 (SENT_SEM: 0.86, SD=0.35; SENT_PERC: 0.73, SD=0.45, PIC_SEM: 0.89, SD=0.31, PIC_PERC: 0.79, SD=0.41). Accuracy was higher for the semantic task than for the perceptual task ($\beta$=0.88, SE=0.09, $p<.001$) and lower for sentence stimuli compared to picture stimuli ($\beta$=-0.28, SE=0.09, $p<.001$), with no interaction between stimulus type and task.

*Experiment 2.* Average response rate was 0.89. Overall mean reaction time was 1.51s, SD=0.49 (SENT_SEM: 1.41s, SD=0.56; SENT_PERC: 1.62s, SD=0.44, PIC_SEM: 1.43s, SD=0.42, PIC_PERC: 1.57s, SD=0.49). RTs for the semantic task were slightly lower than for the perceptual task ($\beta$=-0.17, SE=0.02, $p<.001$), with an interaction between stimulus type and task ($\beta$=-0.08, SE=0.03, $p$=0.016). There was no main effect of stimulus type on reaction time. Overall mean accuracy was 0.77, SD=0.42 (SENT_SEM: 0.64, SD=0.48; SENT_PERC: 0.84, SD=0.36, PIC_SEM: 0.77, SD=0.42, PIC_PERC: 0.84, SD=0.37). Accuracy was lower for the semantic task than for the perceptual task ($\beta$=-0.79, SE=0.09, $p<.001$) and for sentence stimuli compared to picture stimuli ($\beta$=-0.33, SE=0.09, $p<.001$), with an interaction between stimulus type and task ($\beta$=-0.75, SE=0.19, $p<.001$).

*Experiment 3.* Average response rate was 0.90. Overall mean reaction time was 1.09s, SD=0.52 (SENT_SEM: 1.12s, SD=0.6; SENT_PERC: 1.03s, SD=0.51, PIC_SEM: 1.14s, SD=0.47, PIC_PERC: 1.06s, SD=0.5). RTs for the semantic task were slightly higher than for the perceptual task ($\beta$=0.09, SE=0.02, $p<.001$). There was no significant effect of stimulus type (sentences vs. pictures) and no interaction between stimulus type and task. Overall mean accuracy was 0.68, SD=0.47 (SENT_SEM: 0.81, SD=0.39; SENT_PERC: 0.78, SD=0.42, PIC_SEM: 0.75, SD=0.43, PIC_PERC: 0.73, SD=0.44). Accuracy was higher for the semantic task than for the perceptual task ($\beta$=0.16, SE=0.07, $p$=0.031) There was no significant effect of stimulus type and no interaction between stimulus type and task. For 16 participants, accuracy data from one run was erased because of a bug in the script.

Overall, there is no consistent trend in difficulty patterns across experiments, suggesting that the observed neural response patterns cannot be explained by between-condition differences in difficulty.

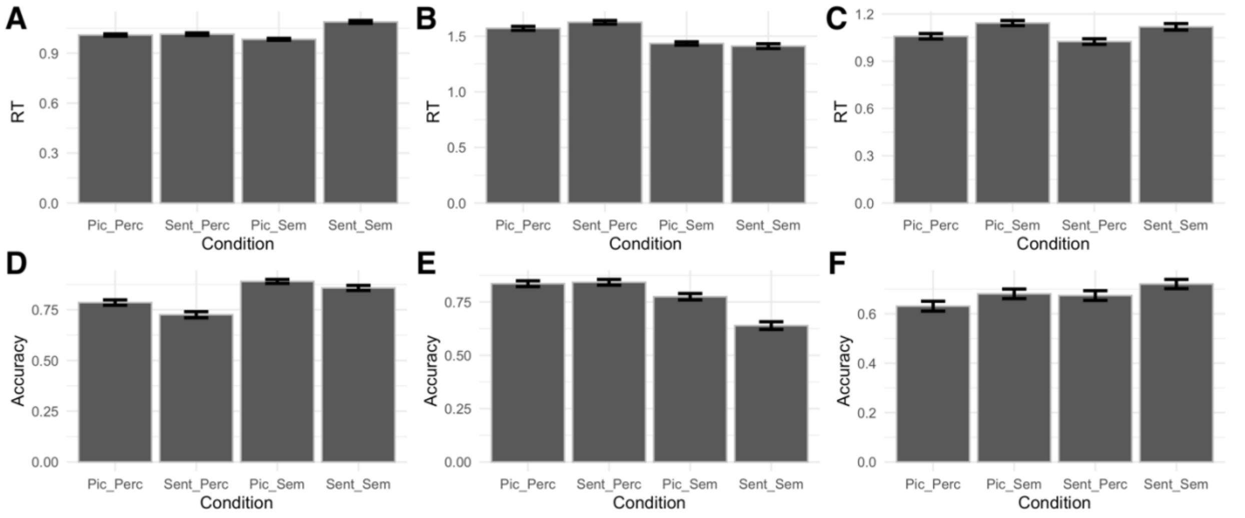


***Figure S1.*** *Behavioral results for the critical experiments.* ***(A-C)*** *Response times for Experiments 1, 2 and 3 respectively.* ***(D-F)*** *Accuracies for Experiments 1, 2, and 3, respectively.*

#### SI-2. Additional information about the responses in semantic fROIs

***Semantic fROI responses generalize across experiments.*** Despite some variability in response magnitudes, overall response patterns are similar across experiments and fROIs (**Figure S2;** see main text for statistics).

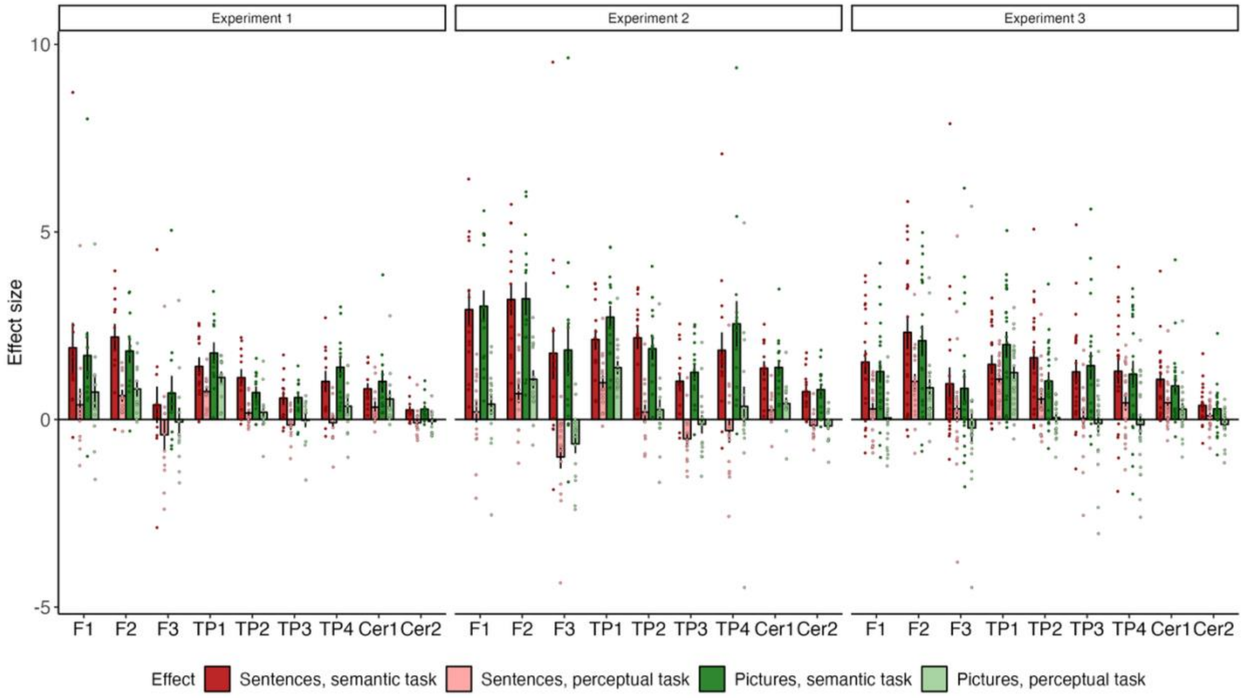


***Figure S2***. *Semantic fROI responses in each experiment.*

***Visual parcels.*** Our whole-brain GcSS analysis yielded 11 parcels, 9 of which are described in the main text. Two parcels appear to be driven by low-level visual properties of the stimuli and are therefore excluded from the main analyses. Their locations and response profiles are shown in **Figure S3**.

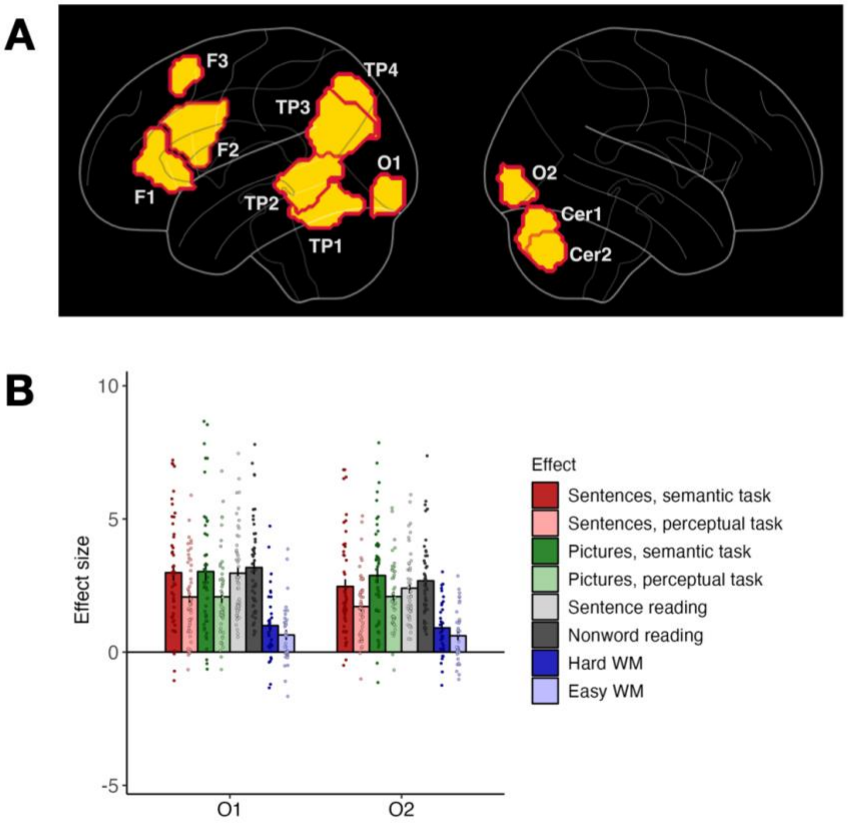


***Figure S3****. fROIs in bilateral early visual cortex were found as part of the brain-whole analysis but excluded from follow-up analyses due to our focus on cognitive rather than perceptual processing.* ***(A)*** *All parcels found via GcSS, including two in the occipital cortex: O1 (left hemisphere) and O2 (right hemisphere).* ***(B)*** *Response profiles of fROIs defined within the occipital parcels.*

***Semantic regions' responses generalize to left-out participants.*** We use independent sets of data (different runs of the same task) when we define a fROI using data from a given experiment and measure its responses to the conditions from the same experiment. In addition, to test how well our results generalize to a new set of participants, we split the participants into two groups: the first group (n=30, 10 from each experiment) were used to define the parcels, and the second group (n=11) was left out. The results were consistent across the two groups: we observed a preference for semantic over perceptual tasks (both groups: $p<.001$), no overall effect of stimulus type, no interaction between stimulus type and task, no preference for sentences over nonwords or for hard vs. easy versions of the working memory task (**Figure S4**).

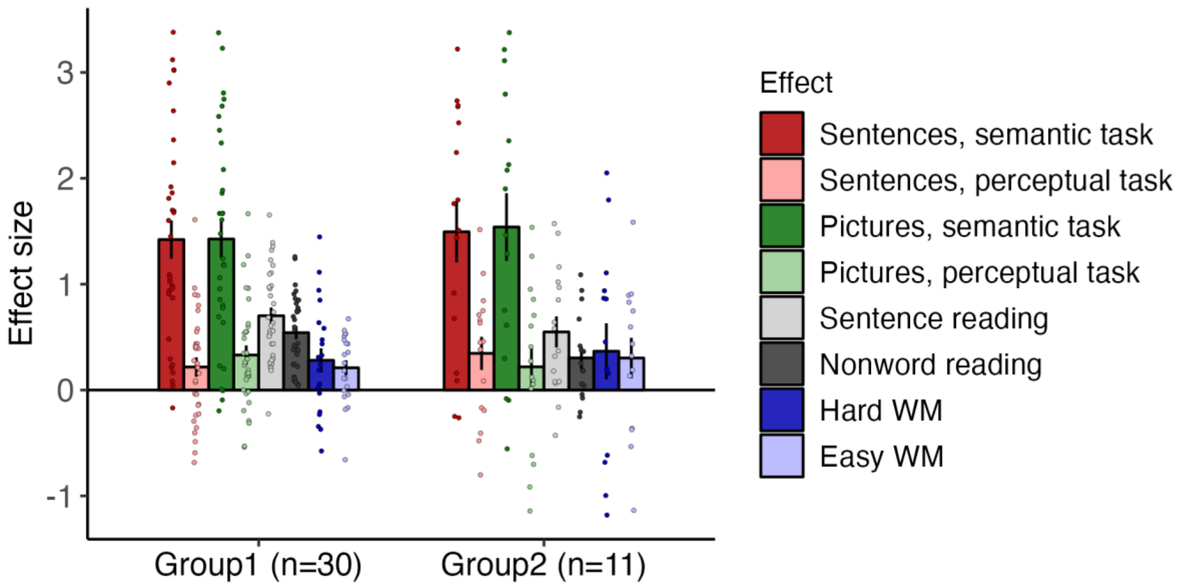


***Figure S4***. *The results generalize to a new group of participants (whose data was not used for parcel definition). WM = working memory.*

***Contralateral semantic fROIs.*** We additionally examined responses in semantic parcels mirrored onto the right cerebral hemisphere and left cerebellum (the original parcels we found were located in the left cerebral hemisphere and right cerebellum). The response profiles of fROIs defined with these contralateral parcels are shown in **Figure S5**; see main text for summary and stats.

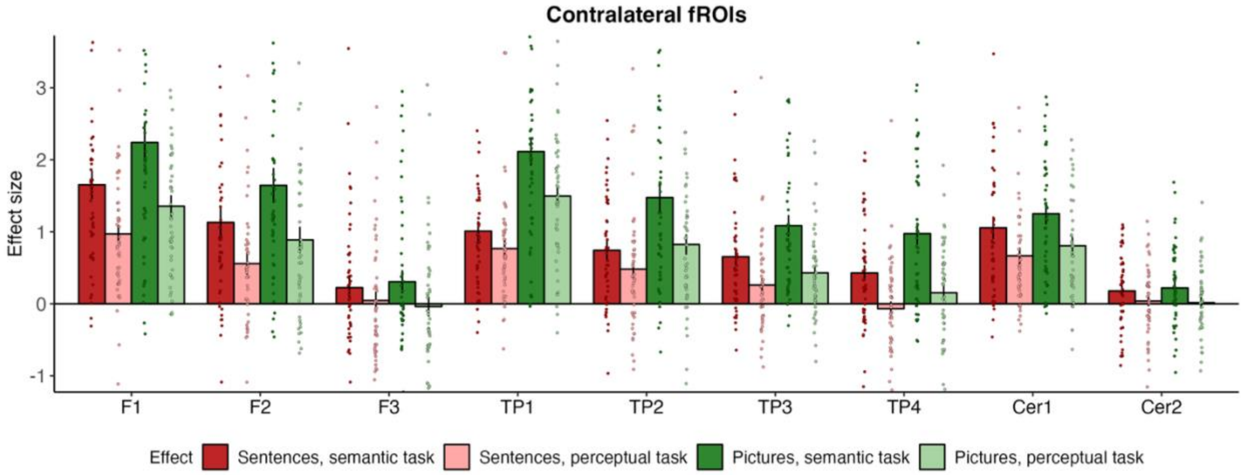


***Figure S5.*** *Response profiles of semantic fROIs defined using contralateral parcels (original semantic parcels projected onto the opposite hemisphere).*

#### SI-4. tSNR analysis details

To assess the temporal stability of the fMRI data, we calculated the temporal signal-to-noise ratio (tSNR) for each voxel across all subjects and functional runs (**Figure S6, A**). For each run, we used preprocessed data that underwent motion correction, spatial smoothing, and normalization to MNI space. We then generated voxel-wise tSNR maps by dividing the mean signal intensity by the temporal standard deviation of the signal at each voxel (Welvaert and Rosseel, 2013) and log transformed them for visualization purposes. To avoid division by zero, voxels with zero standard deviation were assigned a tSNR value of zero. A small constant (1e-10) was added to all tSNR values before log transformation to prevent undefined values.
Next, we calculated parcel-wise tSNR values to examine differences between semantic demand parcels and anatomical anterior temporal lobe (ATL) parcels. Semantic demand parcels were generated using the GcSS approach, while anatomical ATL parcels were adapted based on the Desikan Atlas (Desikan et al., 2006). Specifically, we selected the inf+pole+mid temporal lobe parcels and halved those anatomical regions along the anterior-posterior axis (y-axis) to isolate the anterior portion (**Figure S6, B**).
Using these anatomically defined parcels, we restricted the whole-brain tSNR maps to calculate the mean tSNR within each parcel. This allowed for a direct comparison of tSNR across regions of interest (**Figure S6, C**).

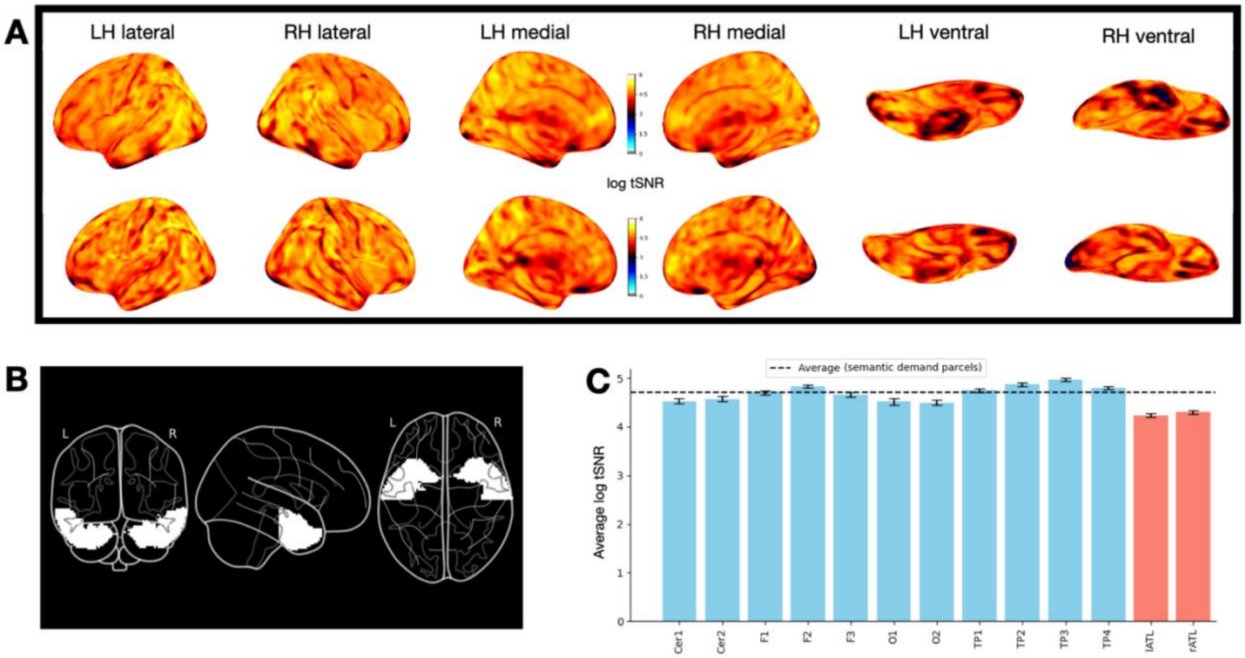


***Figure S6***. *tSNR analysis. (**A**) Sample subject tSNR maps. (**B**) Anatomical ATL parcels. (**C**) Average tSNR in the semantic demand parcels discovered with whole-brain GSS analysis (blue) and in anatomically defined ATL parcels (red).*

#### SI-5. Comparison with previously described semantics-responsive brain regions

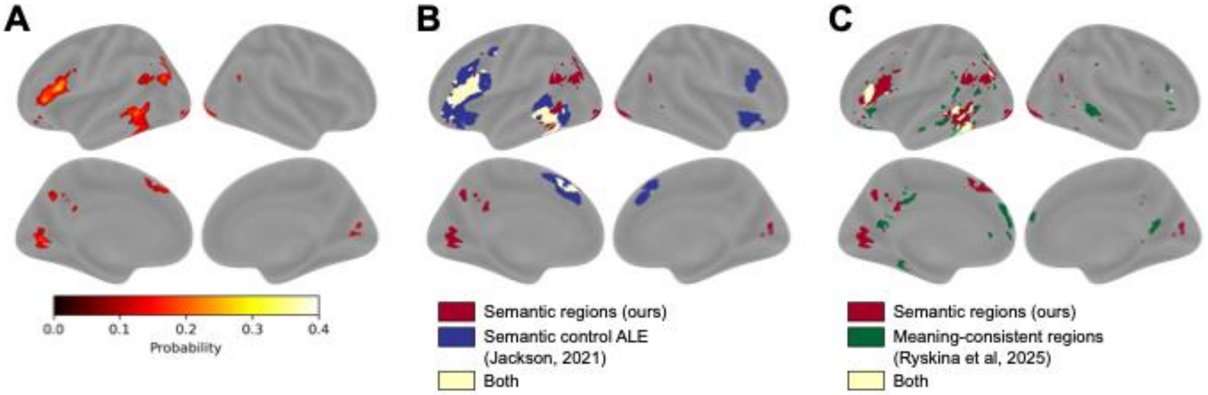


***Figure S7***. *Group-level comparisons of semantic regions' locations.* ***(A)*** *Probabilistic overlap map from which the semantic parcels were derived. Colorbar: ratio of participants who showed significant activation at this voxel.* ***(B)*** *Overlap between semantic regions (ours) and the semantic control regions derived from the meta-analysis in Jackson (2021). ALE = activation likelihood estimation.* ***(C)*** *Overlap between semantic regions (ours) and the meaning-consistent regions from Ryskina et al (2025).*
